# spaCraft: calibrated power analysis and sample-size planning for multi-sample spatial transcriptomics

**DOI:** 10.64898/2026.09.04.749534

**Authors:** Jungmin Shin, Juan Xie, Xiaojie Jin, Qin Ma, Dongjun Chung

## Abstract

Comparative spatial transcriptomics is now routine, yet the number of tissue sections per group is rarely determined by formal power analysis. Power depends jointly on between-sample variation and domains recovered by clustering, a combination not represented by existing tools. We present spaCraft, which converts a replicated pilot into endpoint-specific per-group sample-size recommendations. It fits models of spatial expression, domain geometry and composition, then estimates power through a generate–recover–test loop that re-estimates domains in every synthetic sample, allowing clustering uncertainty to enter the recommendation. Its differential-expression and composition endpoints are tested on recovered rather than assumed domains, yielding calibrated power rather than detection rates. We applied spaCraft to four cohorts spanning Visium, Stereo-seq, and Visium HD. In held-out validation, three-sample pilots predicted sample-size requirements in independent real samples, supporting the full chain from pilot fitting through domain recovery to endpoint testing. Sample size thereby becomes an explicit, reproducible property of the planned analysis rather than an informal guess.

---

Spatial transcriptomics (ST) has been rapidly adopted across tissues and disease contexts, with comparative studies increasingly analyzed at the cohort rather than single-section level ^1,2^. Such studies contrast cases with controls^3^, responders with non-responders, and successive developmental stages^4^. In these designs, the unit of replication is a tissue section, which we refer to as a sample, and the between-condition effect is defined by the contrast across groups rather than by variation within any one sample. The central experimental design question is therefore not how deeply to sequence a single sample, but how many samples each group requires for a prespecified test to detect a between-condition difference with adequate power. Because each ST sample is costly and often irreplaceable ^1,5^, this choice is consequential. Yet sample size is usually determined informally from platform capacity or precedent, leaving its statistical rationale implicit.

Formalizing this decision before data collection requires two linked components, namely a valid hypothesis test and synthetic cohorts that represent the future study at each candidate sample size. The test defines the power function, whereas the synthetic cohorts provide the data on which it is estimated. The first requirement is inferential. Power is the probability that a prespecified test rejects its null under a given alternative, and the power function describes how this probability varies with effect size and sample size. Following the Neyman–Pearson framework^6^, a valid power analysis must therefore be defined with respect to a prespecified hypothesis test, with an explicit null hypothesis, a test statistic with a characterized null distribution, and control of the relevant error criterion under single or multiple testing. Without a valid test and controlled rejection rule, a curve based on whether an effect is detected represents an empirical detection rate rather than a power function and yields no formally interpretable sample-size recommendation.

The second requirement is generative. Because pilot studies rarely contain enough observed samples to estimate power at each candidate size, the required cohorts must be generated from the observed pilot cohort. These synthetic cohorts must preserve the sources of variation that determine power, including between-sample variation in domain composition and geometry, within-sample spatial correlation in expression, and analytical uncertainty from domain recovery. A model fitted to a single pilot sample cannot estimate variation defined across samples, whereas treating recovered labels as fixed removes clustering error. Both simplifications suppress variability and can bias power upward, producing recommendations that may not transfer to the study ultimately conducted. The two requirements are inseparable. Representative synthetic cohorts without valid testing yield detection rates, whereas valid testing without representative cohorts does not estimate power for the planned study. A design framework for comparative ST must therefore connect an observed pilot cohort, generated synthetic cohorts, and valid hypothesis testing within a single procedure.

Each requirement has been addressed in existing work, though largely in separate lines of research. Non-spatial power-analysis frameworks provide valid hypothesis tests and formal sample-size calculations for differential expression and cell-type composition, and are well established for designs in which spatial position plays no role ^7–10^. Spatial simulators reproduce tissue organization and expression patterns with high fidelity and supply realistic data for benchmarking and method development^11–13^. Because they are formulated at the level of a single section, between-sample variation lies outside their scope.

Several approaches move closer to spatial study design, but differ in their replication units and inferential targets. PoweREST^14^ estimates gene-level differential-expression power for Visium by bootstrap resampling spots within prespecified regions, without modeling between-sample variation from a replicated pilot or uncertainty in data-derived domains. The in silico tissue framework (IST)^15^ uses parameterized synthetic tissues to determine how many cells or fields of view are needed to detect cell types and spatial organization, focusing on within-tissue sampling rather than between-condition inference across biological samples. For NanoString GeoMx ^16^, a mixed-effects approach incorporates patient- and region-level replication for investigator-selected regions, but those regions are defined before analysis and the procedure was developed for a specific clinical setting.

Each framework is therefore well matched to the design question for which it was developed, but none combines the elements required for comparative, spot-resolved ST. Table 1 compares their replication units, inferential bases, region definitions, and treatment of between-sample variation. The missing combination is tissue samples as biological replicates, calibrated tests applied to domains recovered during analysis, and between-sample variation estimated from a replicated pilot.

**Table 1.** Design frameworks for spatial power analysis address different configurations. Each row is one configuration, matched to the question the framework was built to answer. Replication unit is the level treated as the independent biological replicate and over which power is indexed. Power basis is the quantity from which the reported power is computed. Region definition is how the regions being compared are obtained, whether fixed before analysis or estimated from the data. Because the frameworks differ on all three, they share no common estimand and a head-to-head benchmark would not be well defined. PoweREST indexes power by the number of tissue samples while drawing its replicate variation from spots resampled within a single region. Comparative, spot-resolved ST calls for the combination in the last row: tissue samples as biological replicates, calibrated tests applied to domains recovered during analysis, and between-sample variation estimated from an observed pilot cohort. The generative performance of spaCraft is compared with spot-resolved simulators in Extended Data Table 1.

| Frame-<br>work | Platform | Replica-<br>tion unit | Inferential | Generative |  |
| --- | --- | --- | --- | --- | --- |
|  |  |  | Power basis | Region defini-<br>tion | Between-sample vari-<br>ation |
| PoweR-EST<sup>14</sup> | Visium | Tissue sam-<br>ples | Gene-level test, multi-<br>plicity adjusted | Fixed before<br>analysis | Bootstrap of<br>spots within the<br>region |
| IST<sup>15</sup> | Cell-resolved<br>imaging | Cells, fields<br>of view | Detection rates, with<br>tests for selected adja-<br>cency analyses | Simulated tissue<br>organization | Not represented |
| GeoMx<sup>16</sup> | NanoString<br>GeoMx | Patients,<br>regions per<br>patient | Mixed-effects test at<br>the patient level | Selected by the<br>investigator | Patient random<br>effect |
| **spaCraft** | **Sequencing-**<br>**based ST** | **Tissue**<br>**samples** | **Cohort-level tests of**<br>**domain expression**<br>**and composition** | **Recovered**<br>**by clustering**<br>**within the**<br>**design loop** | **Estimated**<br>**from a repli-**<br>**cated pilot** |

We therefore propose spaCraft, a pilot-driven framework that formulates sample-size determination for multi-sample ST as a prospective design problem integrated with the downstream analysis pipeline rather than delegated to a separate simulator. From a pilot cohort, spaCraft fits three probabilistic models for synthetic-cohort generation, namely an additive Gaussian-process (GP) model of spatial expression, a Fisher–Gaussian kernel-mixture model (FGKMM) of domain geometry ^17^, and a binomial-logit model of domain composition. Each component is estimated across multiple pilot samples, so that variation in gene expression, tissue organization, and domain abundance enters generation rather than being fixed at the values of a single representative sample. spaCraft then generates synthetic cohorts under a user-specified effect size and estimates power through a generate–recover–test loop (Fig. 1). Within each iteration it withholds the generator labels, re-estimates the spatial domains in every synthetic sample, and applies the endpoint tests to the recovered domains. For this recovery step, spaCraft implements pBANKSY, a pilot-guided domain-recovery algorithm developed here. pBANKSY builds on BANKSY’s spatial feature representation^18^, but augments it with joint pilot–synthetic embedding, pilot-centroid initialization, and one-to-one domain matching so that domain identities remain anchored to the annotated pilot while the generator labels are withheld. Embedding this recovery step within power estimation allows clustering uncertainty to propagate into the sample-size recommendation rather than treating the recovered labels as fixed.

**Figure 1.**
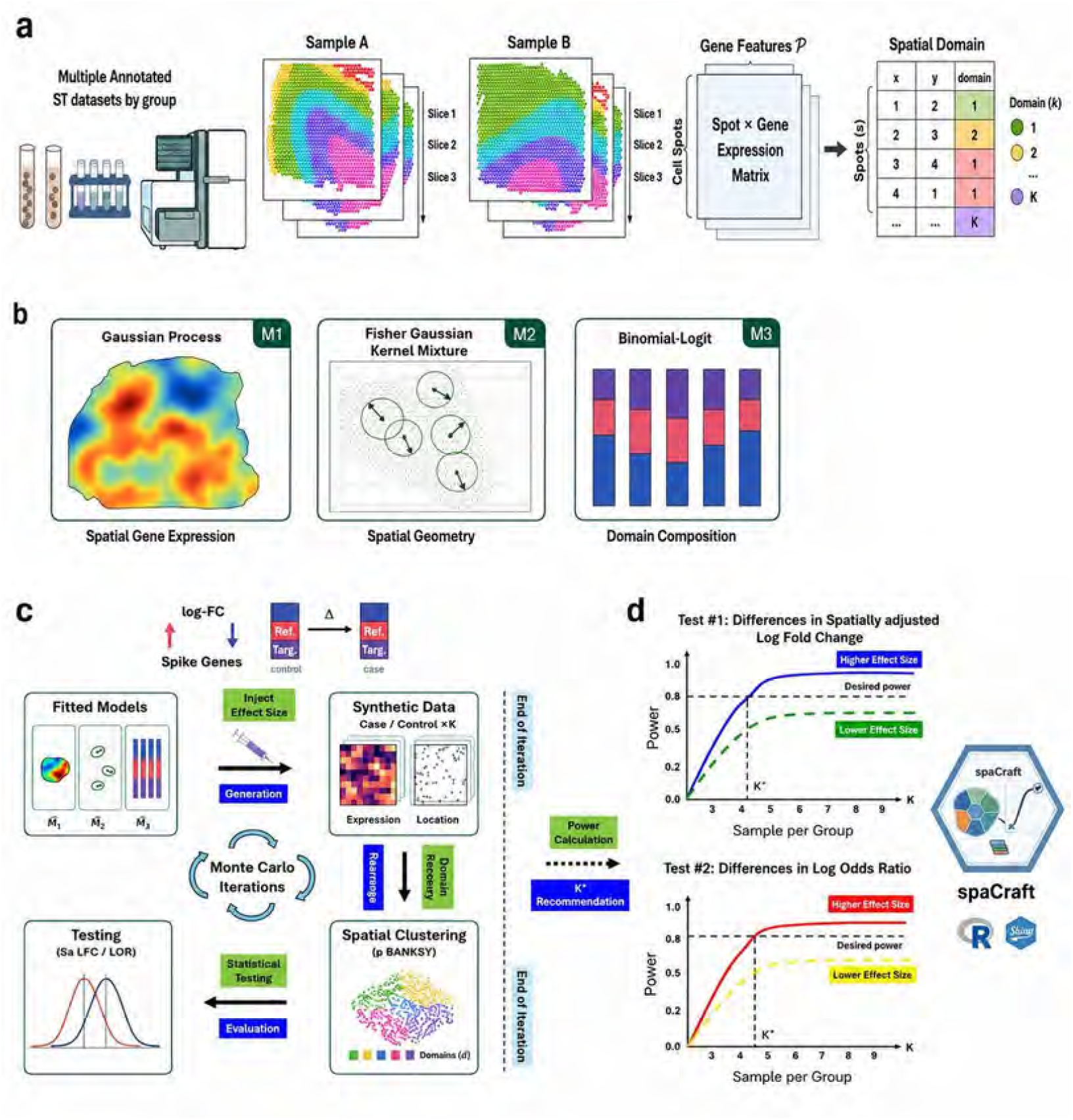
spaCraft turns a pilot cohort into a sample-size recommendation. The workflow is shown as an abstract pipeline, not a specific dataset. **(a)** Input. A group-labeled cohort of annotated ST samples supplies the domain annotation and the spot-by-gene expression matrix. **(b)** Three fitted layers, a Gaussian-process expression model 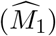, a Fisher–Gaussian kernel-mixture geometry model 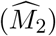, and a binomial-logit composition Model 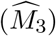. **(c)** The generate–recover–test loop. Each Monte Carlo iteration injects an effect (*p*_target_, *θ*_spike_), generates a synthetic cohort 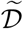 of case and control samples at per-group size *K*, performs domain recovery using pBANKSY, and evaluates SaLFC and LOR on the recovered labels 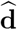. Iterating yields power over *K* and the recommended size 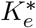. spaCraft is implemented as an R package with a Shiny interface. **(d)** Output. Endpoint power against per-group sample size, SaLFC (top) and LOR (bottom), each at a higher and a lower effect size. 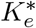 is the smallest *K* reaching the target power *π*^∗^ = 0.8.

Power is reported for two complementary cohort-level endpoints, a spatially adjusted log-fold-change (SaLFC) for differential expression, and a baseline-anchored log-odds-ratio (LOR) for changes in domain composition. The user-specified effect is therefore a target log-fold-change for SaLFC and a target odds ratio for LOR. By default, spaCraft uses Bonferroni-adjusted *P* values for SaLFC, with a Benjamini–Hochberg option for discovery-oriented designs that screen many genes and tolerate a controlled proportion of false discoveries in exchange for greater power.

We applied spaCraft to four cohorts spanning Visium, Stereo-seq, and Visium HD, and these applications yielded two consistent design principles. First, the required cohort size depended on the prespecified endpoint and effect size, and the more demanding endpoint differed across cohorts. Second, the effect relevant to design is the effect retained after domain recovery rather than the effect imposed during generation, because recovery error attenuates the signal reaching the downstream test. Sizing a comparative ST study therefore requires prespecifying both the effect of interest and the analysis through which it will ultimately be tested.

Synthetic fidelity alone, however, does not establish whether a recommendation derived from a small pilot will generalize to future biological samples. We therefore validated this generalizability in an independent Visium mouse brain cohort. Specifically, we repeatedly separated three-sample-per group pilot sets from disjoint validate samples and compared the resulting sample-size requirements. Pilot-based recommendations agreed within one sample with the held-out requirements in 92% of evaluable SaLFC splits and 89% of LOR splits. This agreement provides direct evidence that the full chain from pilot fitting through domain recovery to endpoint testing generalizes beyond the samples used for estimation.

Together, these results show that the inferential and generative requirements of power analysis can be met within a single procedure. By coupling pilot-derived cohort generation, in-loop domain recovery, and calibrated endpoint testing, spaCraft converts a small replicated pilot into an analysis-aware, endpoint-specific sample-size recommendation applicable across sequencing-based ST platforms. It is delivered as an open-source R package with a companion Shiny application. Sample size thereby becomes an explicit, reproducible property of the planned spatial analysis rather than an informal consequence of platform capacity or precedent.

## Results

### spaCraft turns a pilot cohort into a sample-size recommendation

spaCraft converts an observed pilot cohort into endpoint-specific per-group sample-size recommendations. Its inputs are an annotated, group-labeled pilot dataset with spatial domain annotations, with at least two samples per group and a prespecified target–reference domain pair (*T, R*) (Fig. 1a). The dorsolateral prefrontal cortex (DLPFC)^19^ cohort serves as the running example throughout. Because this neurotypical cohort contains no biological case–control contrast, the control and case labels used here denote pseudo-groups defined for workflow illustration (Methods). For notation, we refer readers to the Methods (Table 2).

**Table 2.** Notation. The sample index (*k, c*) is shown when needed and omitted otherwise, as in 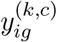 and *D*^(*k,c*)^. An observed cohort contains the samples from both groups, with synthetic and recovered cohorts defined analogously.

| Symbol | Definition |
| --- | --- |
| $c \in \{0, 1\}$ | group index for control (=0) and case (=1) |
| $k = 1, \dots, K_c$ | sample index within group $c$, with $K_0 = K_1 = K$ for balanced designs |
| $i = 1, \dots, N^{(k,c)}$ | spot index in sample $(k, c)$ |
| $g = 1, \dots, G$ | gene index |
| $X_{ig}$ | raw count of gene $g$ at spot $i$ |
| $y_{ig} = \log(1 + X_{ig})$ | log expression used in $M_1$ and SaLFC |
| $\mathbf{s}_i \in \mathbb{R}^2$ | spot coordinate normalized to $[0, 1]^2$ within each sample |
| $\mathcal{A} = \{1, \dots, D\}$ | set of spatial domains |
| $(T, R) \in \mathcal{A}^2, T \neq R$ | prespecified target and reference domains |
| $n_d$ | number of spots assigned to domain $d$ |
| $d_i$ | annotated domain label observed in the pilot for $i$th spot in $(k, c)$ sample |
| $\tilde{d}_i$ | generator label assigned during synthetic-sample generation |
| $\hat{d}_i$ | recovered label estimated by pBANKSY |
| $\mathcal{D} = (\mathbf{X}, \mathbf{S}, \mathbf{d})$ | observed sample triplet |
| $\tilde{\mathcal{D}} = (\tilde{\mathbf{X}}, \tilde{\mathbf{S}}, \tilde{\mathbf{d}})$ | synthetic sample triplet |
| $\hat{\mathcal{D}} = (\tilde{\mathbf{X}}, \tilde{\mathbf{S}}, \hat{\mathbf{d}})$ | recovered sample triplet used for endpoint testing |
| $\mathcal{G}_{\text{SVG}}$ | labeling set of spatially variable genes used for domain recovery |
| $\mathcal{G}_{\text{Null}}$ | empirical-null genes used for SaLFC testing |
| $\mathcal{G}_{\text{spike}} (\subset \mathcal{G}_{\text{Null}})$ | injection set of effect-carrying genes receiving the imposed shift |

The pilot is fitted once using three probabilistic models of spatial expression, domain geometry, and domain composition, together with their variation across samples. These are a Gaussian-process-based additive model of spatial expression (*M*_1_), a Fisher-Gaussian kernel-mixture model (FGKMM) of domain geometry (*M*_2_), and a binomial-logit model of domain composition (*M*_3_) (Fig. 1b). For the expression endpoint, the pilot also defines a labeling set of spatially variable genes (SVGs) for domain recovery and an empirical-null gene set, from which the injection set of effect-carrying genes is selected (Methods).

Each design scenario specifies an effect on an interpretable scale. For expression, *θ*_spike_ is an additive log-fold-change applied to the effect-carrying genes in the case group, with *θ*_spike_ = 0 denoting the null. For composition, *p*_target_ ∈ (0, 1) is the case-group share of the target domain within (*T, R*), with *p*_target_ = *p*_0_ denoting the pilot baseline. The same compositional effect can be expressed as an odds ratio relative to *p*_0_ on the logit scale (Methods).

For each candidate per-group size *K*, spaCraft repeats a generate-recover-test loop over Monte Carlo iterations (Fig. 1c). Each iteration generates a synthetic cohort under the specified effect, withholds the generator labels, and re-estimates the domains with pBANKSY, spaCraft’s pilot-guided domain-recovery algorithm. The coordinates are then rearranged onto the platform lattice (Methods), after which SaLFC and LOR are applied to the recovered labels. Domain-recovery uncertainty therefore enters the same endpoint tests that will be used in the planned analysis.

Repeating this procedure over grids of effect sizes and candidate values of (*K*) yields raw Monte Carlo power estimates, which are smoothed over (*K*) using a monotone shape-constrained additive model ^20^. The fitted curve 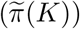 determines the smallest candidate size (*K*^∗^) attaining the target power (*π*^∗^), set to 0.8 by default (Fig. 1d). SaLFC power is the Monte Carlo mean proportion of injected genes declared significant, whereas LOR power is the Monte Carlo rejection probability of the compositional test (Methods). Here and throughout, SaLFC power denotes the expected proportion of prespecified effect-carrying genes declared significant in a future cohort, whereas LOR power denotes the probability that the single domain-composition test rejects its null hypothesis. Because both are valid hypothesis tests applied to recovered domains, the resulting curves estimate endpoint-specific power functions rather than empirical detection rates.

### Expression, geometry, and composition models capture the variation relevant to power

Power in comparative ST depends on variation across biological samples and spatial dependence within each sample. spaCraft represents these sources through three pilot-derived models of spatial expression, domain geometry, and domain composition, each estimating a group-level tissue pattern together with its variation across samples. In the DLPFC analysis, all three models were fitted from three pilot samples per group.

The expression model *M*_1_ decomposes the spot-level log-expression of each gene into sample-specific domain means, a sample-level shift, a spatially correlated GP field, and independent noise. The domain means and sample-level shift represent between-sample expression variation, whereas the GP field and noise represent within-sample spatial and residual variation (Methods). In the DLPFC cohort, 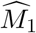 reproduced the laminar expression patterns of representative marker genes (Fig. 2a).

**Figure 2.**
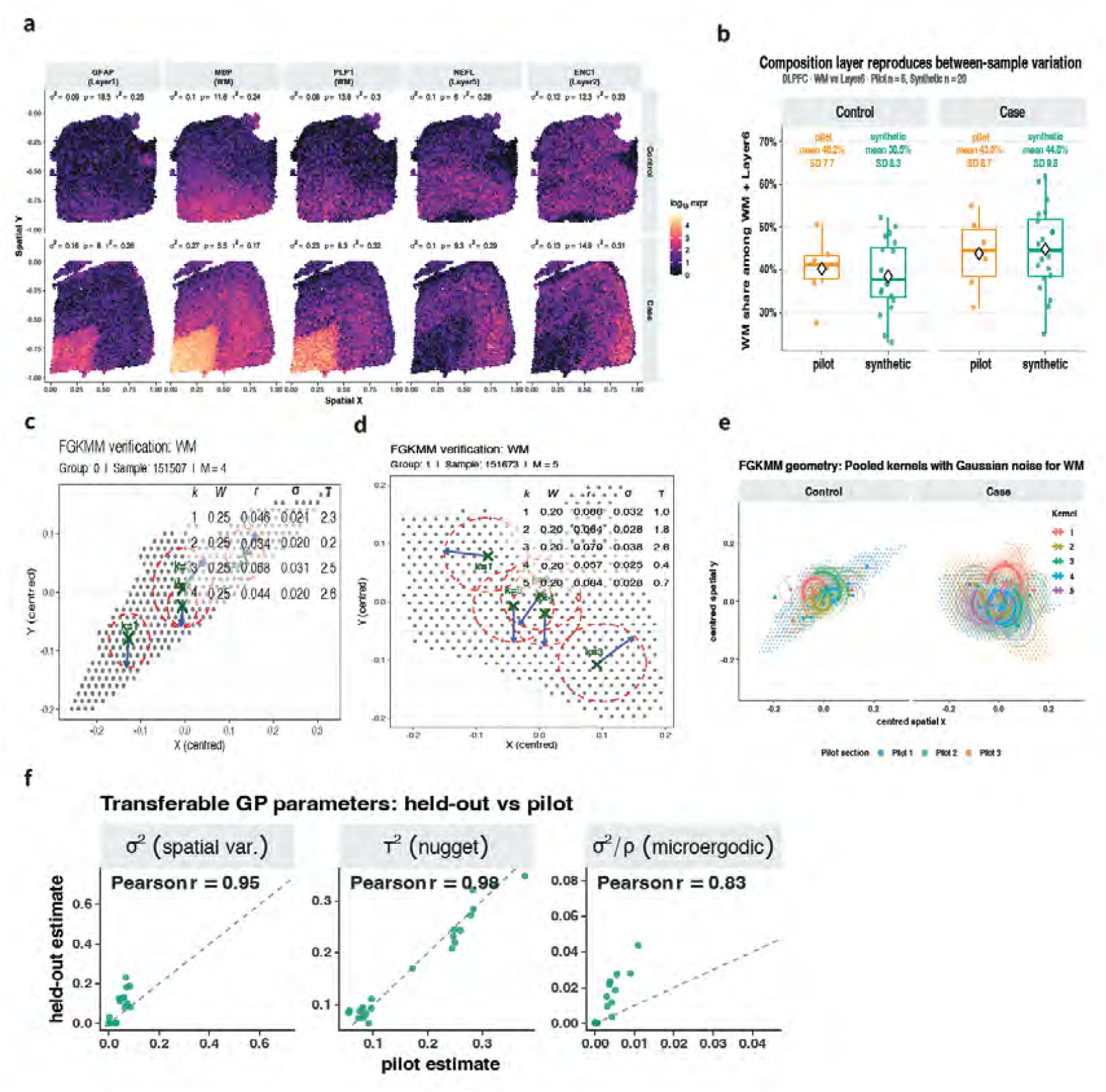
Three pilot-derived models capture variation across DLPFC samples. The target-reference pair is white matter (WM) versus Layer 6. **(a)** Expression model 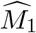. Spatial log-expression of five representative marker genes, *GFAP* for Layer 1, *MBP* and *PLP1* for WM, *NEFL* for Layer 5, and *ENC1* for Layer 2, in control and case pilot samples. The fitted Gaussian-process parameters are the partial sill *σ*^2^, range *ρ*, and nugget *τ* ^2^ for each group. **(b)** Composition model 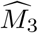. A synthetic cohort (*n* = 20) reproduces the mean and sample-to-sample spread of the WM share observed in the pilot (*n* = 6) within both groups. Diamonds indicate group means. **(c,d)** Geometry model 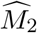 fitted to individual samples. FGKMM representations of the WM domain are shown for one control sample (**c**, *M* = 4 kernels) and one case sample (**d**, *M* = 5 kernels). The displayed kernel parameters are the weight *W*, radius *r*, radial dispersion *σ*, and angular concentration *τ* . These parameters are specific to the geometry model and are distinct from the Gaussian-process parameters in **a. (e)** Pooled geometry model 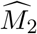. Kernels pooled across pilot samples represent variation in WM position, orientation, and spread. **(f)** Agreement of expression-model parameters with held-out samples. Per-gene estimates from the fitting samples are compared with estimates from held-out samples of the same group. The partial sill *σ*^2^ (*r* = 0.95), nugget *τ* ^2^ (*r* = 0.98), and microergodic ratio *σ*^2^*/ρ* (*r* = 0.83) show agreement across samples. Under fixed-domain asymptotics, *σ*^2^*/ρ*, rather than the range *ρ* alone, is identifiable.

The fitted covariance structure also generalized to samples not used for estimation. Across genes, estimates from the fitting and non-fitting samples of the same group showed Pearson correlations of *r* = 0.95 for the partial sill 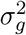 and *r* = 0.98 for the nugget 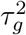 (Fig. 2f). The correlation range *ρ*_*g*_ showed little agreement on its own, whereas the microergodic ratio 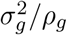, the only quantity consistently estimable under fixed-domain asymptotics^21^, showed a correlation of *r* = 0.83 (Methods and Supplementary Note S2).

Testing is confined to three gene sets fixed from the pilot. The labeling set of spatially variable genes is used for recovery, the empirical-null set provides genes without a group difference, and the injection set is drawn from the null genes with the largest baseline target–reference separation. Because misassignment perturbs these genes most strongly, power is evaluated under a conservative rather than an optimistic scenario (Methods).

The geometry model *M*_2_ separates domain placement from shape, representing domain centroids through their across-sample distribution and domain shape through an FGKMM, which accommodates irregular, non-convex, and disconnected domains that a single parametric shape cannot represent (Methods). In the DLPFC pilot, 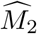 reconstructed the white-matter geometry of individual samples (Fig. 2c,d), and kernels pooled across pilot samples further captured variation in domain position, orientation, and spread (Fig. 2e).

The composition model *M*_3_ represents the target-domain share within the pair (*T, R*) using a binomial-logit model, with its group mean and between-sample dispersion estimated from the pilot (Methods). It reproduced the group means closely, with synthetic versus observed shares of 38.5% versus 40.2% in controls (s.d. 8.3 versus 7.7 percentage points) and 44.8% versus 43.8% in cases (s.d. 9.8 versus 8.7 percentage points) (Fig. 2b).

Similar agreement was observed in an independent mouse-brain cohort from the 5xFAD transgenic model of Alzheimer’s disease^3^, using hippocampus and cortex as the target– reference pair (Extended Data Fig. 1a–d), showing that the three-model fit was not limited to the DLPFC example.

### Synthetic samples reproduce the pilot and carry the specified effects

Each Monte Carlo iteration generates a synthetic sample as 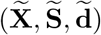, where 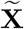 and 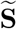 are drawn from the fitted models 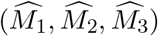 and 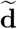 contains generator-assigned domain labels. These labels inherit the pilot domain identities and are allocated according to the generated geometry and specified composition rather than estimated by clustering.

We evaluated two generator-level properties. Pilot fidelity measures how closely the synthetic data reproduce the observed pilot, whereas generator-level effect realization measures whether each specified effect is imposed at its intended size under 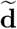. We first assessed pilot fidelity. Within the DLPFC reference sample used for model fitting, the synthetic expression maps faithfully recapitulated the original spatial patterns (Extended Data Fig. 2a). Furthermore, the summary statistics of the synthetic and reference data demonstrated near-perfect agreement across all genes, yielding Pearson correlations of *r* = 1.00 for mean expression, *r* = 0.94 for variance, *r* = 0.99 for detection frequency, *r* = 0.98 for spatial autocorrelation, and *r* = 0.95 for gene–gene correlation (Extended Data Fig. 2b). The dedicated single-sample simulators SRTsim and scDesign3^12^ showed comparable overall fidelity. scDesign3 reproduced per-gene variance more closely, and both simulators better preserved gene-gene correlation (Extended Data Table 1). Because spaCraft generates genes independently given the domain labels, the generated data retain the correlation induced by shared domain membership but not the residual co-expression within domains, a component that does not enter the SaLFC power (Methods and Supplementary Note S5). Pilot fidelity is therefore a necessary property of the generator but is not, by itself, sufficient to validate a cohort-size recommendation.

We next assessed effect realization. Across the scenario grid, the realized effects under 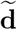 closely followed their specified values, with the expression effect widening the target– reference separation and the compositional effect increasing the target-domain share (Extended Data Fig. 3a,b and Supplementary Fig. S1). Each effect was therefore imposed on its intended scale before domain recovery. Because the endpoint tests use the recovered labels 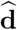 rather than 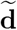, any discrepancy arising after recovery is attributable to the recovery step, which we examine next.

**Figure 3.**
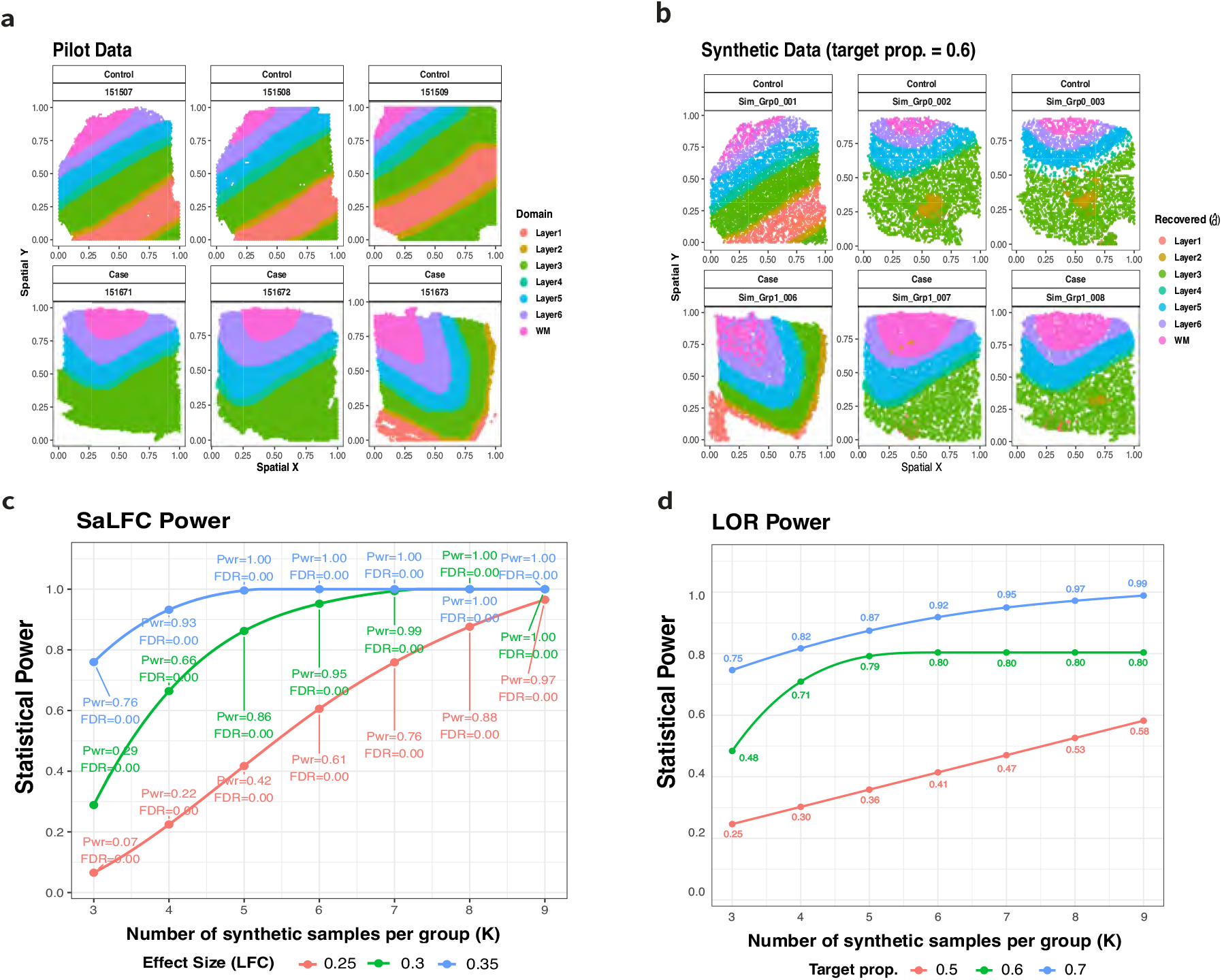
Recovered-domain testing yields endpoint-specific power curves and sample-size recommendations. DLPFC, with white matter (WM) and Layer 6 as the target-reference pair. **(a)** Annotated domains in the six pilot samples, comprising three control and three case samples across Layers 1-6 and WM. **(b)** Domains recovered by pBANKSY in synthetic control and case samples generated with a case target share of 0.6. The recovered domains are displayed after coordinate rearrangement onto the platform lattice (Methods and Supplementary Note S8). **(c)** Monotone-smoothed SaLFC power 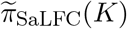 across per-group sample sizes *K* for specified log-fold-changes of 0.25, 0.30, and 0.35. Curves are fitted to raw Monte Carlo estimates, defined as the mean proportion of injected genes declared significant using Bonferroni-adjusted *P* ≤ 0.05 (Methods). Labels show power and empirical FDR. **(d)** Monotone-smoothed LOR power 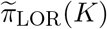 across *K* for specified case target shares of 0.5, 0.6, and 0.7. Curves are fitted to raw Monte Carlo rejection probabilities from the compositional test applied to each recovered cohort (Methods).

### In-loop domain recovery propagates clustering uncertainty into effect fidelity and power

Most domain-level ST analyses test on labels estimated by clustering, so their statistics inherit clustering uncertainty in addition to biological variation across samples. A sample-size calculation should account for both sources.

spaCraft incorporates the planned recovery procedure directly into power estimation. Each Monte Carlo iteration withholds the generator labels 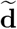 and re-estimates the domains, and endpoint testing uses only the recovered labels 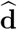, so the estimated power reflects the clustering uncertainty that the downstream analysis will encounter. Recovery must return the same domain identities in every iteration, because the target–reference contrast is defined on named domains rather than on arbitrary cluster indices. In spaCraft, this step is carried out by pBANKSY, the framework’s pilot-guided domain-recovery algorithm. It anchors each synthetic sample to the annotated pilot through a joint embedding, initialization at the pilot-domain centroids, and one-to-one identity matching (Methods). Pilot annotations guide the recovery but are not copied to synthetic spots, and 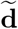 remains hidden, so a spot with 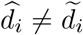 constitutes a misassignment, and the discrepancy between 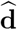 and 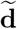 represents domain-recovery error.

We define analysis-aware effect fidelity as the extent to which a specified effect is transmitted through domain recovery to the estimand evaluated by the test statistics. It is quantified by comparing the effect realized under 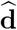 with both its specified value and its value under 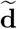.

We benchmarked pBANKSY as the recovery algorithm within the spaCraft loop against the spatial methods BayesSpace^22^ and SpaGCN^23^ and the non-spatial method Seurat^24^ by substituting each into the same recovery step. Among the evaluated procedures, pBANKSY provided the best balance between effect-recovery accuracy and runtime (Extended Data Fig. 4a), the highest design-efficiency index (Extended Data Fig. 4b), and the greatest run-to-run reproducibility (Extended Data Fig. 4c). Runtime and peak memory remained practical on Visium HD, the largest platform evaluated, with more than 10^5^ bins per sample (Extended Data Fig. 5a,b).

**Figure 4.**
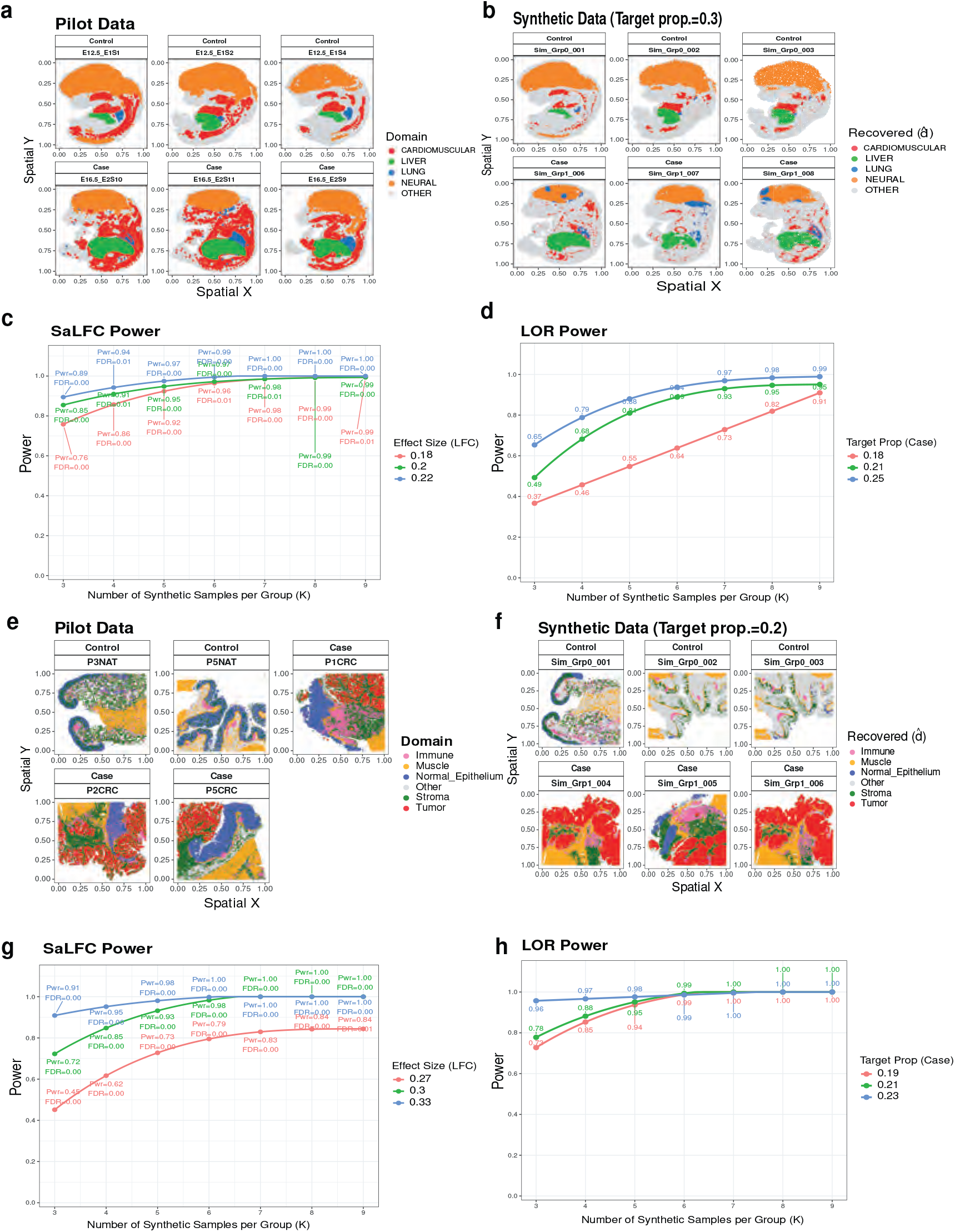
The spaCraft design workflow transfers across sequencing-based ST platforms. **(a-d)** Stereo-seq mouse embryos comparing E12.5 and E16.5, with lung and neural tissue as the target-reference pair. **(e-h)** Visium HD colorectal cancer comparing normal-adjacent and tumor samples, with normal epithelium and stroma as the target-reference pair. **(a,e)** Annotated pilot domains. **(b,f)** Domains recovered by pBANKSY in synthetic samples and displayed after coordinate rearrangement onto the platform lattice (Methods and Supplementary Note S8). **(c,g)** Monotone-smoothed SaLFC power 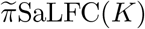 across per-group sample sizes *K* and three specified log-fold-changes. Labels show fitted power and empirical FDR. **(d,h)** Monotone-smoothed LOR power 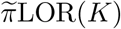 across *K* and three specified case target shares. Curves are fitted to the corresponding Monte Carlo estimates (Methods).

**Figure 5.**
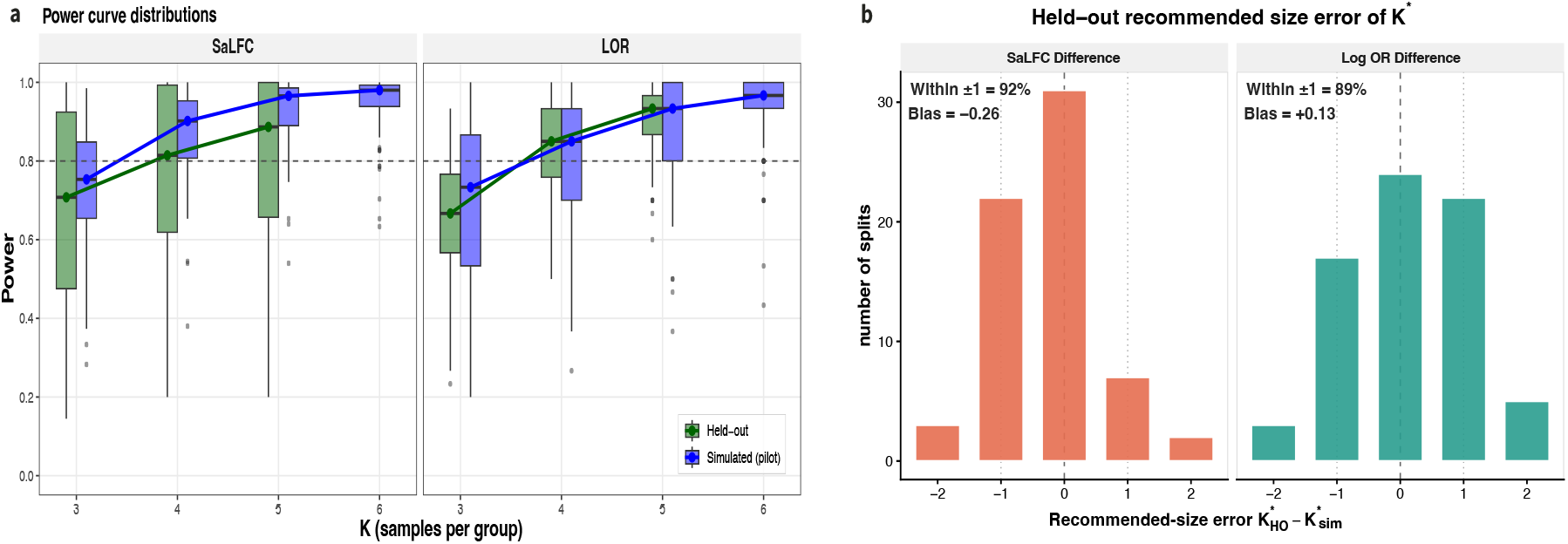
Pilot-driven recommendations agree with held-out real samples. The 5xFAD Visium cohort 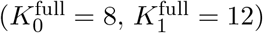, hippocampus versus cortex, is split *B* = 100 times into a pilot of three samples per group and its disjoint complement. Both sides run the full generate–recover–test loop and are evaluated on pBANKSY-recovered domains. **(a)** Pilot-simulated and held-out power for both endpoints over all *B* = 100 splits, with the dashed line marking the target *π*^∗^ = 0.8. The held-out grid stops at *K* = 5, the largest balanced size its smaller arm admits, and the two distributions overlap within their interquartile ranges at every size both sides admit. **(b)** Recommended-size error 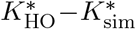 in samples, with zero denoting exact agreement, over the splits on which *K*^∗^ is identified on both sides and lies within the range the held-out arms can express (*n* = 65 for SaLFC, *n* = 71 for LOR). The error is within one sample in 92% and 89% of splits, at mean −0.26 and +0.13.

In the DLPFC analysis, pBANKSY recovered structured spatial domains while leaving residual misassignments across cortical layers (Fig. 3a,b). These recovery errors translated into different analysis-aware effect fidelity for the two endpoints (Extended Data Fig. 3c). The realized log-fold-change under 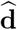 remained close to its value under 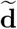 and to its specified value, because averaging over many spots within each recovered domain absorbs a small fraction of mislabeled spots. The realized odds ratio dropped from 4.9 under 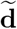 to 2.3 under 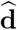 against a specified 5, because misassigned spots enter the target and reference counts directly. In both cases recovery acted toward the null, and no spurious compositional difference arose under the compositional null. Power is therefore calculated from the effect retained under 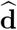, so this attenuation is carried into the sample-size recommendation.

### Endpoint-specific power curves yield sample-size recommendations

We next converted the effects retained under the recovered labels 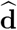 into two endpoint-specific power curves. For each candidate per-group size *K* and specified effect, we applied the SaLFC test with Bonferroni-adjusted *P* values and the single compositional LOR test to the prespecified target–reference pair (Methods). The software also provides the Wilcoxon rank-sum test as an optional nonparametric alternative, although all analyses reported here use SaLFC and LOR.

The monotone-smoothed SaLFC power increased with both *K* and the specified log-fold-change (Fig. 3c). At an effect size of 0.35, the fitted power was 0.93 at *K* = 4 and 1.00 at *K* = 5. At 0.30, the fitted curve first exceeded the target power of 0.8 at *K* = 5 (0.86), whereas an effect of 0.25 required *K* = 8 (0.88). The fitted LOR power likewise increased with *K* and the specified case target share (Fig. 3d). At a target share of 0.7, the fitted curve first exceeded 0.8 at *K* = 4 (0.82). At 0.6, it first reached the target at *K* = 6, whereas at 0.5, the specified share closest to the control baseline, fitted power remained below 0.6 even at *K* = 9.

The endpoint tests underlying these fitted power curves remained calibrated. In the evaluated SaLFC settings, the empirical false discovery rate remained below the nominal level of the Bonferroni-adjusted procedure, and under the compositional null the empirical type I error of the LOR test remained near the nominal level across the evaluated values of *K* (Supplementary Fig. S2 and Methods).

For endpoint *e*, spaCraft defines *K*_*e*_ as the smallest candidate per-group size at which the monotone-smoothed power curve 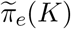 reaches the target power *π*. The recommendation is therefore specific to both the endpoint and the assumed effect. In the DLPFC analysis, reducing the target log-fold-change from 0.30 to 0.25 increased 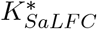 from five to eight, whereas LOR required four samples per group for a target share of 0.7 but did not reach the target within the evaluated range for 0.5. A design adequate for one endpoint or effect size may therefore be inadequate for another. The same pattern was observed in the 5xFAD analysis (Extended Data Fig. 6a–d). Each recommendation is reported with its endpoint, effect size, target power, and error criterion.

These four elements are design decisions that precede data collection, and the minimal adequate cohort is the smallest *K* consistent with all of them jointly. spaCraft converts each such plan into its power curve and 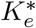. Because both endpoints reduce to two-sample comparisons of per-sample summaries, any endpoint admitting such a summary can enter the same loop once its effect scale, test, and rejection rule are prespecified. More broadly, spaCraft provides a general foundation for extensible spatial study design, allowing user-defined endpoints to be incorporated within the same generate–recover–test framework.

### spaCraft supports study design across sequencing-based ST platforms

The spaCraft workflow transferred across sequencing-based ST platforms with different spatial resolutions and data structures, without modification to its modeling, recovery, or testing procedures. We applied the same workflow to a Stereo-seq mouse-embryo cohort comparing embryonic day 12.5 (E12.5) with embryonic day 16.5 (E16.5) ^4^, and a Visium HD colorectal-cancer cohort comparing normal-adjacent and tumor samples^25^. The Visium HD cohort also contained a non-convex tumor-stroma interface, providing a distinct test of the geometry model. Only platform-specific preprocessing was required (Methods). The target–reference pairs were lung versus neural tissue for Stereo-seq (Fig. 4a-d), and normal epithelium versus stroma for Visium HD (Fig. 4e-h).

For each application, we fitted the same three models (Supplementary Figs. S3 and S4), selected gene sets using the same pilot-based selection criteria (Supplementary Table S4), and applied the same generate–recover–test loop. The synthetic cohorts reproduced the overall spatial organization of both pilots, including the irregular tumor-stroma boundary in the Visium HD samples (Fig. 4b,f).

Both endpoints yielded monotone power curves across the two platforms (Fig. 4c,d,g,h), and the empirical FDR of SaLFC remained below *α* under Bonferroni-adjusted testing in all evaluated settings. The required sample size remained endpoint-specific. In the embryo analysis, 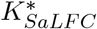 ranged from three to four samples per group across the evaluated effects, whereas 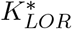 ranged from five to eight. In the tumor analysis, the smallest evaluated effects required four samples per group for LOR and seven for SaLFC. The ordering of the two requirements thus reversed between cohorts, so the more demanding endpoint is determined by the tissue, the contrast, and the effect size rather than by the endpoint itself.

### spaCraft recommendations agree with held-out sample-size requirements

Having established that the pilot-fitted model components transferred to non-fitting samples at the parameter level (Fig. 2f), we asked whether this transfer persisted through the full workflow. The central validation question was whether a recommendation learned from a small replicated pilot predicted the sample size required by independent samples from the same cohort.

We evaluated this transfer using a Visium mouse-brain cohort from the 5xFAD model, comprising 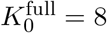 control and 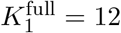 case samples, with hippocampus as the target domain *T* and cortex as the reference *R*. This cohort provided sufficient replication in both groups for repeated disjoint pilot and held-out partitions.

We generated *B* = 100 group-stratified partitions, each containing a pilot of three samples per group and a disjoint held-out set of five control and nine case samples. All models were fitted using the pilot alone. For each partition, the full generate–recover–test loop produced a pilot-simulated power curve and recommendation 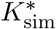. The same specified effect was then applied to the held-out samples, their domains were recovered using the same pBANKSY procedure, and the corresponding requirement 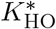 was determined at *π*^∗^ = 0.8 (Methods). The comparison evaluated the full analysis-aware workflow, including biological heterogeneity across samples and analytical uncertainty from domain recovery.

The simulated and held-out power estimates were concordant across partitions (Fig. 5a). At the candidate sizes supported by both analyses, *K* ∈ {3, 4, 5}, their interquartile ranges overlapped for both endpoints, although the held-out distributions were wider. The median difference *π*^HO^ − *π*^sim^ remained within 0.034 of zero.

The resulting sample-size recommendations also agreed (Fig. 5b). Because the smaller held-out arm contained five samples, held-out power could be evaluated only through *K* = 5. Partitions in which either power curve did not reach the target within this range were therefore not included in the *K* agreement summary. Among the remaining evaluable partitions, 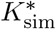 was within one sample of 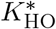 in 92% of SaLFC partitions and 89% of LOR partitions. The mean signed difference 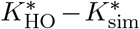 was −0.26 samples for SaLFC and +0.13 samples for LOR, and both were statistically equivalent to zero under a margin of one sample (*p <* 10^−9^). The variation of 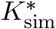 across pilot subsets further quantified sensitivity to pilot-sample selection. Details are reported in Supplementary Table S1, Supplementary Note S10, and Supplementary Table S2.

## Discussion

We developed spaCraft, a pilot-driven framework for prospective cohort design in comparative spatial transcriptomics. It integrates the inferential and generative requirements of power analysis by fitting pilot-derived models of spatial expression, domain geometry, and domain composition, then generating cohorts, recovering their domains, and applying endpoint tests to the recovered labels. The resulting recommended sample size is therefore specific to the planned endpoint, effect size, target power, error criterion, and recovery procedure.

Across applications, two findings had direct implications for prospective study design. First, a universally optimal sample size cannot be defined independently of the planned analysis. The minimum adequate cohort depends on the endpoint and effect being targeted, and when several endpoints must all be adequately powered, the largest requirement determines the cohort. Rather than seeking a single study-wide sample size, spaCraft formalizes these dependencies and provides a statistically principled, biologically interpretable, and accessible framework for identifying the minimum sample size consistent with a pre-specified design. Second, the relevant effect for design is the effect retained after domain recovery. Because recovery error can attenuate the signal available to downstream testing, prospective design should account for the effect that survives the planned analysis rather than the effect imposed during generation.

These design implications are useful only if the underlying sample-size recommendations are reliable. Three complementary results support this. First, the power curves were derived from analysis-aware, calibrated hypothesis tests rather than empirical detection frequencies. The empirical FDR of the default SaLFC procedure remained below 0.05 under Bonferroni-adjusted testing, and the null rejection rate of LOR remained near its nominal level. Second, the same modeling, recovery, and testing procedures produced endpoint-specific power curves across Visium, Stereo-seq, and Visium HD, requiring only platform-specific preprocessing. Finally, recommendations from three-sample-per-group pilots agreed within one sample with requirements measured on independent samples from the same 5xFAD cohort in 92% and 89% of evaluable partitions. This held-out agreement provides evidence that the analysis-aware design procedure can transfer pilot-derived sample-size requirements to independent biological samples, beyond reproducing the synthetic data themselves.

The current study also defines two boundaries. First, the generator draws genes independently conditional on domain labels. This does not affect mean gene-level SaLFC power or Bonferroni validity, but newly defined endpoints that depend on gene–gene correlation, such as co-expression modules or multigene signatures, would require an expression model that preserves residual co-expression, potentially through copula or latent-factor structures. Second, the current implementation targets sequencing-based, spot-resolved data, a two-group contrast, and one target–reference pair at a time. Extensions to imaging-based point patterns, multiple contrasts, or longitudinal designs with repeated samples per subject will require corresponding spatial models, multiplicity control, and tests that account for within-subject dependence.

Every comparative ST study faces an unavoidable sample-size decision, yet that decision has largely rested on informal guesses from platform capacity, budget, or precedent. spaCraft turns this decision into a statistically rigorous, explainable, and reproducible calculation within a unified framework. In this sense, spaCraft follows an intrinsic-hoc strategy by incorporating experimental-design considerations and spatial domain constraints directly into the computational framework^26^. Sample size thereby becomes a property of the planned spatial analysis rather than a consequence of platform capacity. spaCraft is available as an open-source R package (github.com/c16267/spaCraft) with a companion Shiny application (chunglab.bmi.osumc.edu/spaCraft/) and a reference catalog of 21 human and mouse ST datasets spanning Visium, Stereo-seq, and Visium HD. When no pilot is available, investigators can select from this catalog a public cohort matching their tissue, platform, and contrast and use it as a surrogate pilot to obtain a preliminary sample-size estimate before data collection. Laboratories can also upload and analyze their own pilot cohorts.

## Methods

### Notation and gene sets

Table 2 summarizes the notation used throughout.

The distinction among the three domain labels is central to the framework. The annotated labels **d** are observed in the pilot and used for model fitting. The generator assigns 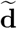 when a synthetic sample is drawn, whereas pBANKSY estimates 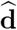 from the synthetic expression and coordinates. Thus, 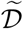 and 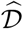 share the same expression and coordinates and differ only in their domain labels. Endpoint testing uses 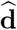, thereby incorporating the domain-recovery error expected in a future analysis.

The three gene sets are selected once from the pilot and held fixed across Monte Carlo replicates. The labeling set *G*_SVG_ of spatially variable genes (SVGs) is used for domain recovery. The empirical-null set *G*_Null_ is used for SaLFC testing, and its subset, the injection set *G*_spike_, receives the imposed expression effect. Because the injected and noninjected genes are selected from the same empirical-null population before generation, they differ only by the imposed shift. Rejections in *G*_spike_ are therefore counted as true positives, and rejections in *G*_Null_ \ *G*_spike_ as false positives. Expression is generated for *G*_SVG_ ∪ *G*_Null_. The specific criteria for selecting each gene set are detailed in Supplementary Note S1, and the resulting sets are listed in Supplementary Tables S3 and S4.

The selection of G_spike_ from the null genes with the largest baseline target–reference separation makes the power estimate conservative. Spot misassignment displaces the per-sample summary of a gene in proportion to its baseline separation 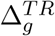 and inflates its between-sample variance accordingly, whereas a gene with 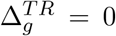 would yield power close to that under perfect recovery. The injection set therefore comprises the members of *G*_Null_ most affected by domain-recovery error (Supplementary Note S1).

### Spatial gene expression model (*M*_1_)

For gene *g* in sample (*k, c*), log expression is modeled as

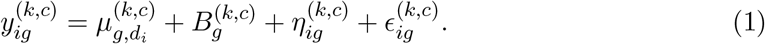

Here, 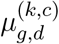 is the sample- and domain-specific mean, 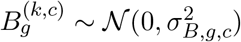 is a sample-level shift shared across domains, 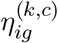 is a mean-zero GP, and 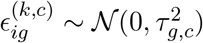 is independent spot-level noise, the three being mutually independent conditional on the domain means and labels. The GP posits isotropic exponential covariance

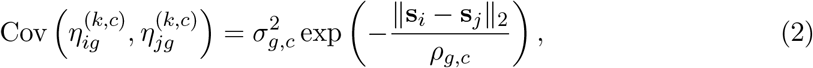

Where 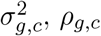, and 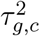 denote the partial sill, range, and nugget variance.

Between-sample variation enters through the sample-level shift 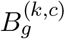 and the sample-specific domain means, with 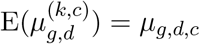 and 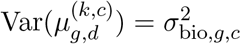 for each domain *d*, and, for the prespecified target and reference domains, 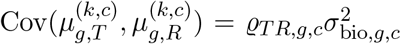. Because 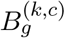 is shared across domains, it cancels from the target–reference contrast.

The model is fitted separately to each pilot sample and pooled within group, all parameters being estimated separately by group. The group mean *µ*_*g,d,c*_ averages the sample-specific domain means, whereas 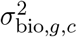 is estimated from their between-sample variation after the shared sample-level shift is removed. Spatial parameters are estimated by maximum likelihood under the nearest-neighbor GP approximation implemented in BRISC^27,28^, which replaces the dense covariance factorizations of the exact likelihood with conditional distributions given small ordered neighbor sets. Group-level parameters are obtained by moment-based pooling,

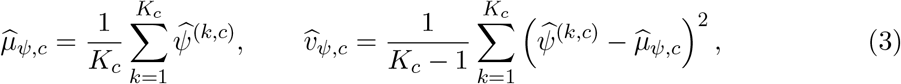

with the analogous empirical covariance for vector-valued parameters. Parameter-specific pooling rules, the joint covariance of the target and reference means, and spatial-parameter identifiability are detailed in Supplementary Note S2.

Equation (1) is specified marginally for each gene, so residual cross-gene covariance is not parameterized. Correlation is nevertheless induced in generated data, because all genes share the same domain labels, geometry, and sample-level structure, and genes with similar domain means covary through 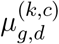. This omission affects neither the SaLFC estimand, an average of marginal gene-level rejection probabilities that dependence leaves unchanged in expectation, nor the default Bonferroni adjustment, which controls the family-wise error rate under arbitrary dependence. It can affect domain recovery, since correlated labeling genes carry less independent information than uncorrelated ones, and the held-out validation bounds the net effect of this and other approximations on the final recommendation.

### Spatial geometry model (*M*_2_)

The geometry model separates domain placement from domain shape, allowing both to vary across samples. The centroid of domain *d* in group *c* follows

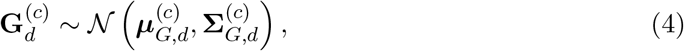

where 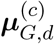 and 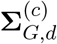 are the empirical mean and covariance of the sample-specific centroids.

Thus, 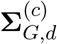 captures between-sample variation in domain position.

Conditional on the centroid, the centered coordinates 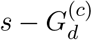 follow an FGKMM^17^.

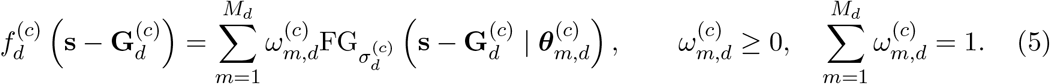

Each component has parameters 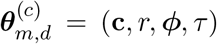 . The pair (**c**, *r*) determines a local center and radius, whereas (***ϕ***, *τ* ) determines direction and angular concentration. The shared parameter 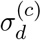 controls radial dispersion. Within *M*_2_, *τ* is therefore an angular-concentration parameter and 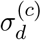 is a geometric dispersion parameter. They are distinct from the nugget variance 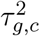 and GP partial sill 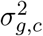 in *M*_1_.

For each group-domain pair, the Bayesian information criterion (BIC) selects the number of components from a candidate set, *M* ∈ {3, …, 6} in the reported analyses. All pilot samples are then refitted by the expectation-maximization (EM) algorithm at the selected *M*_*d*_, keeping the model order fixed across samples. The resulting sample-specific centroids and kernel parameters are combined within group by moment-based pooling. Empirical means and variances or covariances summarize Euclidean parameters, circular moments summarize directions, and matched moments calibrate the scalar parameters and mixture weights. These pooled quantities define group-specific generative priors that preserve between-sample variation in domain position, orientation, and shape. The FG density, EM fitting, parameter-specific pooling, and prior calibration are detailed in Supplementary Note S3.

### Domain composition model (*M*_3_)

The composition model describes the abundance of the target domain relative to a prespecified reference domain while allocating the remaining spot budget separately. For sample (*k, c*), let 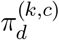 denote the underlying probability that a spot belongs to domain *d* ∈ *A*, with 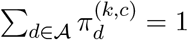, let 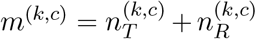 denote the number of spots assigned to the target and reference pair, and let *p*^(*k,c*)^ denote the target share within this pair. Then

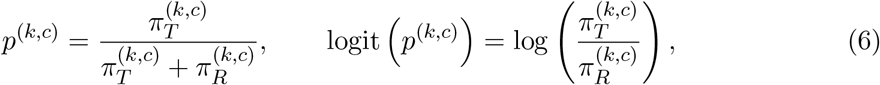

so *p*^(*k,c*)^ is the conditional probability of the target domain given membership in the pair, and the target–reference contrast is invariant to the abundances of the remaining domains. The target count and group-level share are modeled as

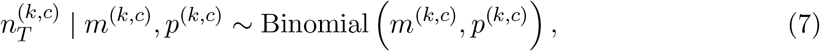

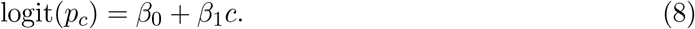

The control and case target shares are *p*_0_ = logit^−1^(*β*_0_) and *p*_1_ = logit^−1^(*β*_0_ + *β*_1_), and the corresponding case-to-control OR is exp(*β*_1_). The coefficients (*β*_0_, *β*_1_) are estimated by a single Binomial generalized linear model fitted jointly to all pilot samples using their observed target counts and pair sizes.

To capture heterogeneity beyond Binomial sampling variation, synthetic samples are allowed to deviate from their group-level target share through a Gaussian random effect on the logit scale. Its dispersion is calibrated from the observed between-sample variation in pilot target shares, and the intercept is adjusted numerically so that *p*_*c*_ remains the marginal mean target share within group *c*.

The remaining composition parameters are estimated by moment-based pooling. Within each group, the target-reference pair budget is summarized by the sample mean and variance of {*m*^(*k,c*)^}, and the background-domain probabilities are obtained by averaging and renormalizing the per-sample residual-domain proportions.

A differential-abundance scenario is parameterized by the marginal case target share *p*_target_ ∈ (0, 1). Given the fitted control baseline *p*_0_, its implied OR is

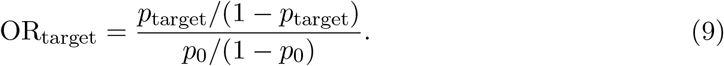

The compositional null is *p*_target_ = *p*_0_, equivalently OR_target_ = 1. When *p*_target_ is not specified, it defaults to the fitted pilot case share *p*_1_. Random-effect calibration, pair-budget pooling, and background-domain allocation are detailed in Supplementary Note S4.

### Synthetic data generation

The three fitted models define probability distributions from which each synthetic sample is drawn,

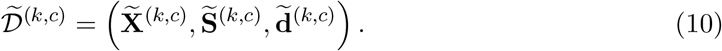

Generation is designed to achieve two objectives. Pilot fidelity preserves the principal expression, geometry, and composition features of the pilot while allowing stochastic variation across samples. Effect realization ensures that the user-specified case-group effects are imposed on their intended scale under the generator labels 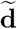.

An effect scenario is specified by the marginal case target share *p*_target_ and the expression shift *θ*_spike_ ≥ 0. Generation proceeds as 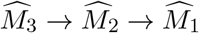, because domain counts determine the number of spots assigned to each domain, and the resulting coordinates determine the spatial expression field.

For each sample, 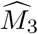 draws the target-reference and background domain counts from the fitted composition model. Control samples retain the fitted baseline share *p*_0_, whereas case samples have marginal target share *p*_target_. Conditional on these counts, 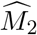 draws the domain centroids, FGKMM parameters, and continuous coordinates, assigning a generator label 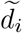 to each spot. Finally, 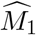 draws the sample-specific domain means, sample-level shifts, spatial effects, and spot-level noise. For *g* ∈ *G*_spike_, *θ*_spike_ is added only to the case target-domain mean. Thus, the two scenario parameters alter the case group while the remaining pilot-derived group and domain structure is retained.

Unrestricted draws from the fitted distributions could occasionally produce spatial patterns that are statistically plausible but poorly representative of the pilot tissue. Generation therefore combines parametric sampling with pilot-anchored components. In 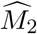, selected FGKMM parameters, including component centers and directions, are blended with a randomly selected pilot cloud to preserve observed layout and orientation. In 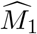, a newly generated parametric spatial field is combined with pilot-conditioned residual texture,

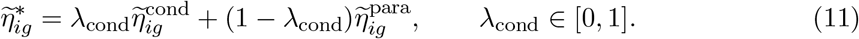

The blended field is centered and rescaled to the fitted partial sill 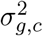 before independent noise with variance 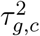 is added. These hybrid constructions preserve pilot-specific geometry and local expression texture without eliminating between-sample stochastic variation. The resulting log expression is transformed to counts by 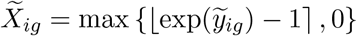.

Genes are generated independently across genes given the generator labels. All genes in a sample share the same label field 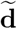, so the generated data reproduce the gene-gene correlation that arises when two genes track the same domains. Residual co-expression within domains, namely the correlation that remains between two genes after conditioning on the labels, is not modeled. This choice is aligned with the primary estimand. SaLFC power is the mean proportion of injected genes declared significant, and by linearity of expectation it depends only on each gene’s marginal rejection probability, namely the probability that the test of that gene alone rejects. The joint behavior of the tests affects only the Monte Carlo variance of the rejected proportion, and the default Bonferroni adjustment controls the family-wise error rate under arbitrary dependence (Supplementary Note S5, section ‘Gene-wise conditional independence and the SaLFC estimand’).

The generated expression and continuous coordinates are then passed to pBANKSY without using the generator labels. Domain recovery replaces 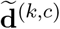 with 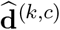, yielding

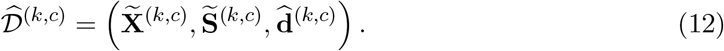

Coordinate rearrangement is performed after the recovered labels have been fixed. The hybrid FGKMM sampling, scalable GP approximation, pilot-conditioned texture, and variance matching are detailed in Supplementary Note S5 (Algorithm S1).

### Domain recovery within the Monte Carlo loop

Domain recovery is an explicit component of the spaCraft Monte Carlo loop and is implemented by pBANKSY, a spaCraft-specific pilot-guided recovery algorithm. pBANKSY adopts BANKSY’s spatial feature construction^18^ but introduces the pilot anchoring required for repeated design simulation: pilot and synthetic spots are embedded jointly, clustering is initialized at annotated pilot-domain centroids, and recovered clusters are assigned pilot domain identities by one-to-one matching. The generator labels 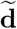 are never used during recovery. For each synthetic sample, pBANKSY maps the generated expression and coordinates 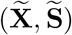 to recovered domain labels 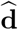. Endpoint testing therefore reflects domains recoverable from the observed data and propagates domain-recovery uncertainty into the estimated power.

For its spatial feature representation, pBANKSY uses the three BANKSY-derived blocks of standardized expression **C**, neighborhood mean **M**, and local dispersion **G**^18^. For the pilot and synthetic sample, respectively,

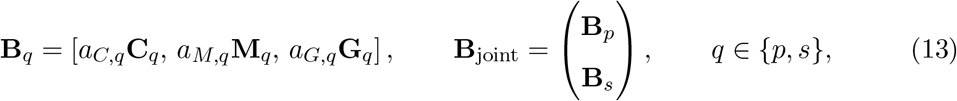

where the scaling weights allocate a fraction *λ*_*p*_ of the feature energy to spatial-neighborhood information.

A single principal component analysis embeds the pilot and synthetic spots in the same feature space. Synthetic spots are then clustered by *k*-means initialized at the annotated pilot-domain centroids. Because cluster indices are arbitrary, the recovered synthetic clusters are assigned pilot domain identities by minimizing

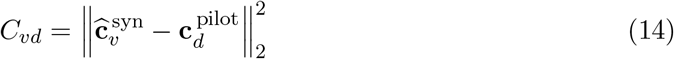

over one-to-one cluster-domain matchings using the Hungarian algorithm. This matching changes only cluster identities, not the recovered partition, and does not use the generator labels. The resulting analysis-ready sample is 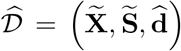, which preserves the pilot domain set and the target-reference interpretation while incorporating realistic recovery error. Feature construction, normalization, tuning of *λ*_*p*_, and centroid matching are detailed in Supplementary Note S6 (Algorithm S2).

Because recovery is repeated across sample sizes, effect scenarios, and Monte Carlo replicates, the clustering method must balance effect-recovery accuracy, computational cost, and run-to-run stability, and this criterion motivated the benchmark reported in Extended Data Fig. 4.

### Weighting scheme

The recovery weight *λ*_*p*_ and texture weight *λ*_cond_ are tuned sequentially from the pilot. First, repeated pilot cross-validation selects *λ*_*p*_ to maximize the mean adjusted Rand index (ARI) between recovered domains and withheld pilot annotations. Conditional on the selected *λ*_*p*_, *λ*_cond_ is then tuned separately for each endpoint and effect scenario.

For each candidate *λ*, the second stage maximizes the regularized noncentrality score

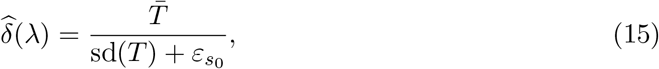

where 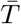 and sd(*T* ) are the empirical mean and standard deviation of the endpoint statistic across tuning replicates. The stabilizing term 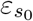 prevents unstable selection when the empirical variation is small. A null guard excludes weights that compromise calibration, so the selected value favors a strong and reproducible effect signal without inflating null rejection. Candidate grids and the complete tuning procedure are given in Supplementary Note S7 (Algorithm S3 and Supplementary Fig. S5).

### Coordinate rearrangement

Because *M*_2_ generates continuous coordinates, a post-recovery rearrangement maps each synthetic point cloud onto the fixed lattice of a pilot sample drawn at random from the same group. Within each recovered domain, a Gaussian optimal-transport map matches the first two spatial moments of the synthetic cloud to those of the corresponding pilot lattice sites,

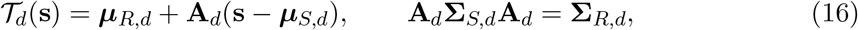

after which the transported points are assigned bijectively to the available sites. When *p*_target_ changes the relative sizes of the target and reference domains, their pooled lattice sites *P* are ordered by *s*(**p**) = *d*(**p, P**_*T*_ ) − *d*(**p, P**_*R*_), and the first *m* = ⌊*p*_target_|*P*|⌋, where ⌊·⌋ denotes the nearest integer function, sites define the target territory. The imposed abundance change is therefore expressed as a coherent boundary shift rather than as isolated relabeled spots.

Rearrangement leaves expression values, recovered labels, domain counts, and the target-reference effect estimate unchanged. It can nevertheless alter the SaLFC test statistic and *P* value because the within-sample covariance depends on pairwise distances between the rearranged coordinates. The LOR statistic is unchanged because it depends only on domain counts. Transport, boundary construction, and lattice-site assignment are detailed in Supplementary Note S8 (Algorithm S4 and Supplementary Fig. S6).

### Endpoint I: Spatially adjusted log-fold-change test (SaLFC)

The SaLFC endpoint tests whether the between-group change in a target-reference expression contrast exceeds a practically meaningful margin. Each biological sample contributes one contrast, preserving the sample rather than the individual spot as the unit of replication. For gene *g*, the recovered labels 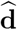 define

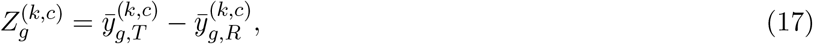

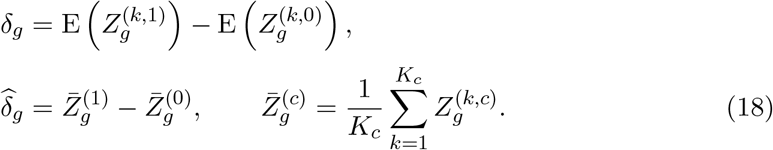

Here, 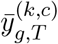 and 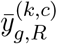 are the mean log-expression values in the recovered target and reference domains. Thus, 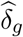 estimates the between-group difference in the corresponding within-sample contrasts.

Its standard error combines spatial uncertainty within tissue sections and biological heterogeneity across samples,

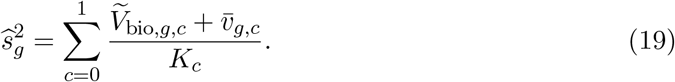

Here, 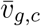 is the group-average variance of the target-reference contrast computed from the fitted GP covariance, and 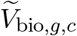 is its empirical-Bayes-moderated between-sample variance^29^. This construction accounts for spatially correlated spots without treating them as independent biological replicates.

Following the Testing Relative to a Threshold (TREAT) principle ^30^, the default analysis uses a right-sided interval test,

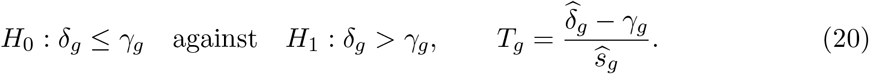

The margin *γ*_*g*_ ≥ 0 defines changes regarded as practically null. By default, it is calibrated as the median magnitude of the gene-specific contrast under null bootstrap resampling, 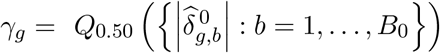. Rejection therefore requires a group difference larger than the fluctuation typically observed under the null, rather than merely a nonzero estimate.

The statistic is evaluated using a *t* distribution with Satterthwaite degrees of freedom. Reverse-direction and two-sided interval alternatives are defined analogously. The GP-based variance calculation, empirical-Bayes moderation, bootstrap calibration, multiplicity adjustment, and asymptotic validity are detailed in Supplementary Note S9.

### Endpoint II: Baseline-anchored compositional test (LOR)

The LOR endpoint tests whether the between-group change in the target-reference composition exceeds both the pilot baseline and a practically meaningful margin. Each biological sample contributes one log-odds contrast, preserving the sample as the unit of replication. For sample (*k, c*), the recovered labels 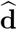 define

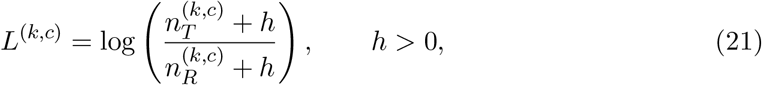

where 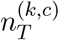 and 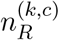 are the recovered target and reference domain counts. The continuity correction *h* keeps the log odds finite when either count is zero.

Let 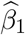 denote the pilot log odds ratio estimated from Eq. (8). Conditional on this fitted baseline, the baseline-anchored effect and its estimator are

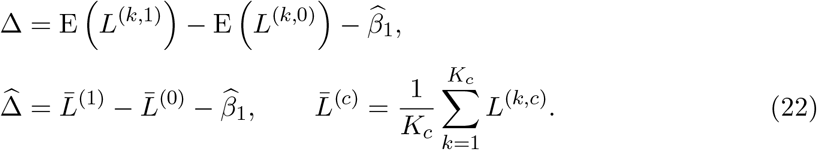

Thus, 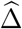 estimates the additional case-to-control compositional change beyond that observed in the pilot. In particular, Δ = 0 corresponds to preservation of the pilot log odds ratio rather than equal target shares between groups.

The standard error allows the between-sample variation in log odds to differ between groups,

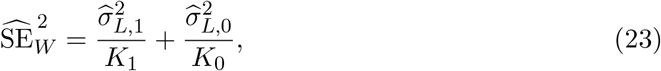

where 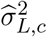 is the empirical variance of 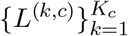 within group *c*. This Welch construction quantifies biological heterogeneity across samples without treating domain counts as independent replicates.

Following the Testing Relative to a Threshold (TREAT) principle ^30^, the default analysis uses a right-sided interval test,

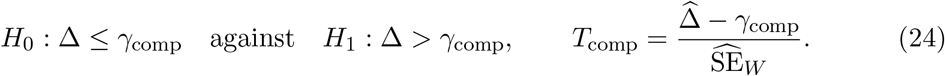

The margin *γ*_comp_ ≥ 0 defines additional compositional changes regarded as practically null. By default, it is calibrated as the median magnitude of the baseline-centered effect under null bootstrap resampling, 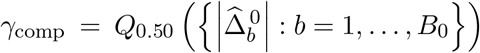. Rejection therefore requires a case-group change beyond the pilot baseline that is also larger than the fluctuation typically observed under the null.

The statistic is evaluated using a *t* distribution with Welch-Satterthwaite degrees of freedom. Reverse-direction and two-sided interval alternatives are defined analogously. Continuity correction, bootstrap calibration, variance estimation, and asymptotic validity are detailed in Supplementary Note S9.

### Power estimation and sample size determination

For each candidate per-group sample size *K* and effect scenario, we generated *N*_rep_ independent synthetic cohorts, recovered their domains with pBANKSY, and applied the corresponding endpoint test at level *α*. Power was estimated from the resulting Monte Carlo rejection decisions. Because SaLFC tests multiple genes within each cohort, whereas LOR performs one compositional test, the two endpoints require distinct power definitions.

For SaLFC, let 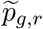 denote the multiplicity-adjusted *P* value for gene *g* in replicate *r*. Bonferroni adjustment is used by default to control the family-wise error rate within each cohort, with the Benjamini-Hochberg procedure available as an option for false discovery rate control. Define 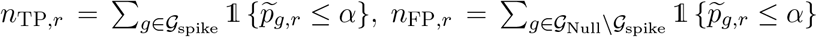, where *n*_rej,*r*_ = *n*_TP,*r*_ + *n*_FP,*r*_. The estimated SaLFC power is the average proportion of injected genes detected,

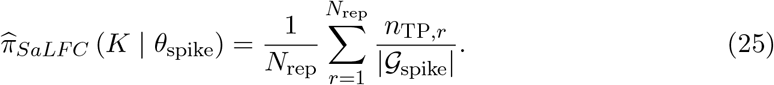

Thus, SaLFC power is the mean gene-level power over *G*_spike_, rather than the probability of detecting at least one or all injected genes. Each SaLFC power curve is presented together with the empirical false discovery rate 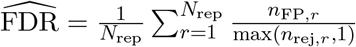. This quantity is the mean false discovery proportion across Monte Carlo replicates under the selected multiplicity adjustment. It serves as an error-calibration diagnostic and does not enter the power estimate or sample-size recommendation. Unless otherwise stated, the reported analyses use Bonferroni adjustment. Because the Bonferroni adjustment controls the family-wise error rate, the empirical FDR is bounded in expectation by the same nominal level, so values below *α* indicate calibration consistent with the procedure’s guarantee.

For LOR, each replicate produces one compositional-test decision, so power is the Monte Carlo rejection probability

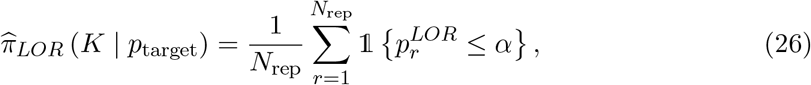

where 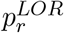 is the LOR *P* value in replicate *r*.

Monte Carlo variation can make the raw power estimates nonmonotone in *K*. For each endpoint and effect scenario, we therefore fit a monotone shape-constrained additive model^20^ to obtain the fitted power curve 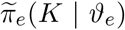, where *ϑ*_*SaLFC*_ = *θ*_spike_ and *ϑ*_*LOR*_ = *p*_target_. For a target power *π*, the recommended per-group sample size is

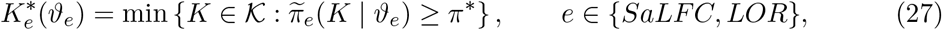

where *K* is the candidate sample-size grid. For SaLFC, *π* is the target expected proportion of injected genes detected under the selected multiplicity adjustment; for LOR, it is the target rejection probability of the single compositional test. We used *π*^∗^ = 0.80 throughout unless otherwise stated. If the target is not reached over *K*, no sample-size recommendation is returned. The power curves shown in the main and Extended Data figures report 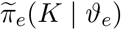.

### Held-out validation procedure

Held-out validation evaluated whether a sample-size recommendation learned from a small pilot transferred to independent real tissue sections. Each cohort was repeatedly partitioned within group into a pilot subset and a disjoint held-out subset. All generative models, gene sets, tuning weights, and bootstrap-calibrated testing margins were estimated from the pilot alone and then held fixed throughout validation.

For each split, the full generate–recover–test procedure produced a pilot-simulated power curve 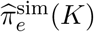 and its recommended per-group sample size 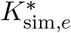 . In parallel, balanced held-out subsamples of *K* samples per group were used to estimate the corresponding held-out power curve 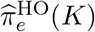 and sample-size requirement 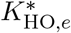 at the same effect size and target power.

For SaLFC, the specified expression shift was added within the annotated case target domain. For LOR, spots were reassigned between the target and reference domains to attain *p*_target_, and expression for reassigned spots was redrawn from 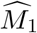 conditional on the new domain. Annotated labels were used only to impose the effect. Both the pilot-simulated and held-out cohorts were subsequently analyzed by pBANKSY, with endpoint testing based only on the recovered labels 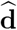.

For retained split *b*, agreement is measured by the recommended-size error

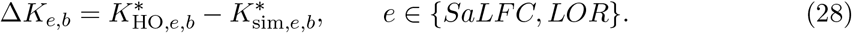

Positive values indicate that the pilot-simulated procedure underestimated the sample size required in held-out samples, whereas negative values indicate a conservative recommendation. Across retained splits, we summarize systematic error and near agreement by

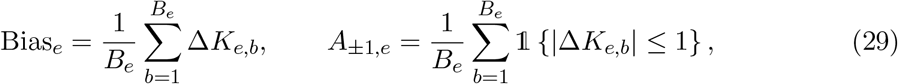

where *B*_*e*_ is the number of splits for which both power curves cross the target power within their evaluated sample-size grids. The distributions of the two power curves and of Δ*K*_*e,b*_ are reported across these splits.

This validation assesses transfer of the final sample-size recommendation, rather than only the visual similarity of synthetic and real data. It therefore evaluates the combined effects of pilot fitting, effect fidelity, domain recovery, endpoint testing, and power-curve estimation. Effect injection, repeated subsampling, grid construction, rounding, exclusion accounting, and sensitivity analyses are detailed in Supplementary Note S10.

### Datasets and preprocessing

Four spatial transcriptomics cohorts were analyzed. Sizes, dimensions, and domain contrasts are in Extended Data Table 2, and domain annotations were taken as released with each study unless stated otherwise.

- **Visium (DLPFC and 5xFAD)**. The DLPFC cohort^19^ comprises 12 sections from three neurotypical donors, with four sections per donor arranged as two pairs of adjacent sections. Because the cohort contains no biological group contrast, we formed two pseudo-groups by sample identifier, control = {151507–151510, 151669, 151670} and case = {151671–151676}, and used samples 151507–151509 and 151671–151673 as the three-per-group pilot. White matter was the target domain and layer 6 the reference. The DLPFC analysis therefore serves to develop and illustrate the spaCraft workflow rather than to represent a biological case–control design. Because multiple sections originate from the same donor, their between-section variation may underestimate the between-subject variation expected in a study of independent biological replicates. The 5xFAD mouse-brain cohort^3^ was used for held-out validation, with hippocampus against cortex.
- **Stereo-seq (mouse embryo)**. Three embryos at embryonic day 12.5 and three at day 16.5^4,31^. Fine-grained annotations were consolidated into five groups, neural, cardiomuscular, liver, lung, and other, and neighboring spots within a group were combined into non-overlapping pseudo-spots by summing counts and averaging coordinates, so that aggregation never merged spots across annotated boundaries. Lung was the target and neural tissue the reference.
- **Visium HD (colorectal cancer)**. Three colorectal adenocarcinoma and two normal-adjacent samples ^25^. We used the 16 *µ*m binned matrices released with the study without further binning, retaining bins with more than 100 counts and more than 50 detected genes. Each sample was normalized, reduced by principal-component analysis, and clustered with Seurat on the first 30 components at resolution 0.5; the sample-specific clusters were then mapped to six common domains, tumor, normal epithelium, stroma, immune, muscle, and other, by canonical marker expression and spatial localization. Normal epithelium was the target and stroma the reference.

All analyses require at least two pilot samples per group, the condition under which the between-sample dispersion components of the three layers are identifiable. A pilot without domain annotations is first clustered and its clusters labeled before fitting.

### Statistics and reproducibility

Unless stated otherwise, the nominal level was *α* = 0.05, the target power was *π*^∗^ = 0.8, and each (*K*, effect) cell used *N*_rep_ = 50 Monte Carlo replicate cohorts. The continuity correction of (21) was *h* = 0.5, and the tuning constants of (15) were *ε* = 10^−5^ with *s*_0_ the median replicate standard deviation of the endpoint statistic over the tuning grid (Supplementary Note S7). Held-out validation used *B* = 100 partitions, *K*_pilot_ = 3, and *N*_rep_ = 30 per side, with splits retained by the prespecified evaluability criterion of Supplementary Note S10. No statistical method was used to predetermine cohort size, no samples were excluded, and randomization and blinding are not applicable because all analyses are simulations based on publicly available datasets.

All Monte Carlo draws were seeded. The master seed is an argument of each top-level entry point of spaCraft 1.0.1, and per-cell seeds are derived deterministically from it across effect sizes, sample sizes, and replicates, so that every power curve and recommendation reported here is reproducible from the seeds recorded in the repository. Analyses were run in R 4.6.1. spaCraft implements pBANKSY internally and uses BRISC 1.0.6 for Gaussian-process fitting, limma 3.68.4 for the empirical-Bayes moderation of (19), and scam 1.2.22 for the monotone shape-constrained smoothing of (27). Seurat 5.5.1 was used for preprocessing and non-spatial clustering, and BayesSpace 1.22.0 and SpaGCN 1.2.7 (Python 3.13.5) entered only the recovery benchmark, in which Seurat also served as the non-spatial comparator. All timings and memory measurements were made on the Ohio Supercomputer Center Ascend cluster using 40 CPU cores.

## Data availability

The DLPFC data are available through the spatialLIBD package (http://spatial.libd.org/spatialLIBD)^19,32^. The 5xFAD data are available from the Gene Expression Omnibus under accession GSE233208^3^. The Stereo-seq MOSTA mouse-embryo data are available from the STOmics database (https://db.cngb.org/stomics/mosta/) under CNGB accession CNP0001543^4,31^. The Visium HD colorectal cancer data are available from the 10x Genomics datasets portal, with raw data in the Gene Expression Omnibus under accession GSE280318^25^. Processed pilot subsets, the simulation files underlying the reported power analyses, and a curated databank of 21 human and mouse spatial transcriptomics datasets are hosted on a publicly accessible laboratory server and can be downloaded through the companion Shiny application (chunglab.bmi.osumc.edu/spaCraft/) or the GitHub repository (github.com/c16267/spaCraft). Source data are provided with this paper.

## Code availability

The spaCraft R package (v1.0.1) is available under the GPL (≥ 3) license at github.com/c16267/spaCraft, together with the seeded scripts that reproduce all power curves and sample-size recommendations reported here. The version used in this study is archived on Zenodo (doi.org/10.5281/zenodo.21907013). A companion Shiny application is available at chunglab.bmi.osumc.edu/spaCraft/.

## Extended Data

**Extended Data Figure 1.**
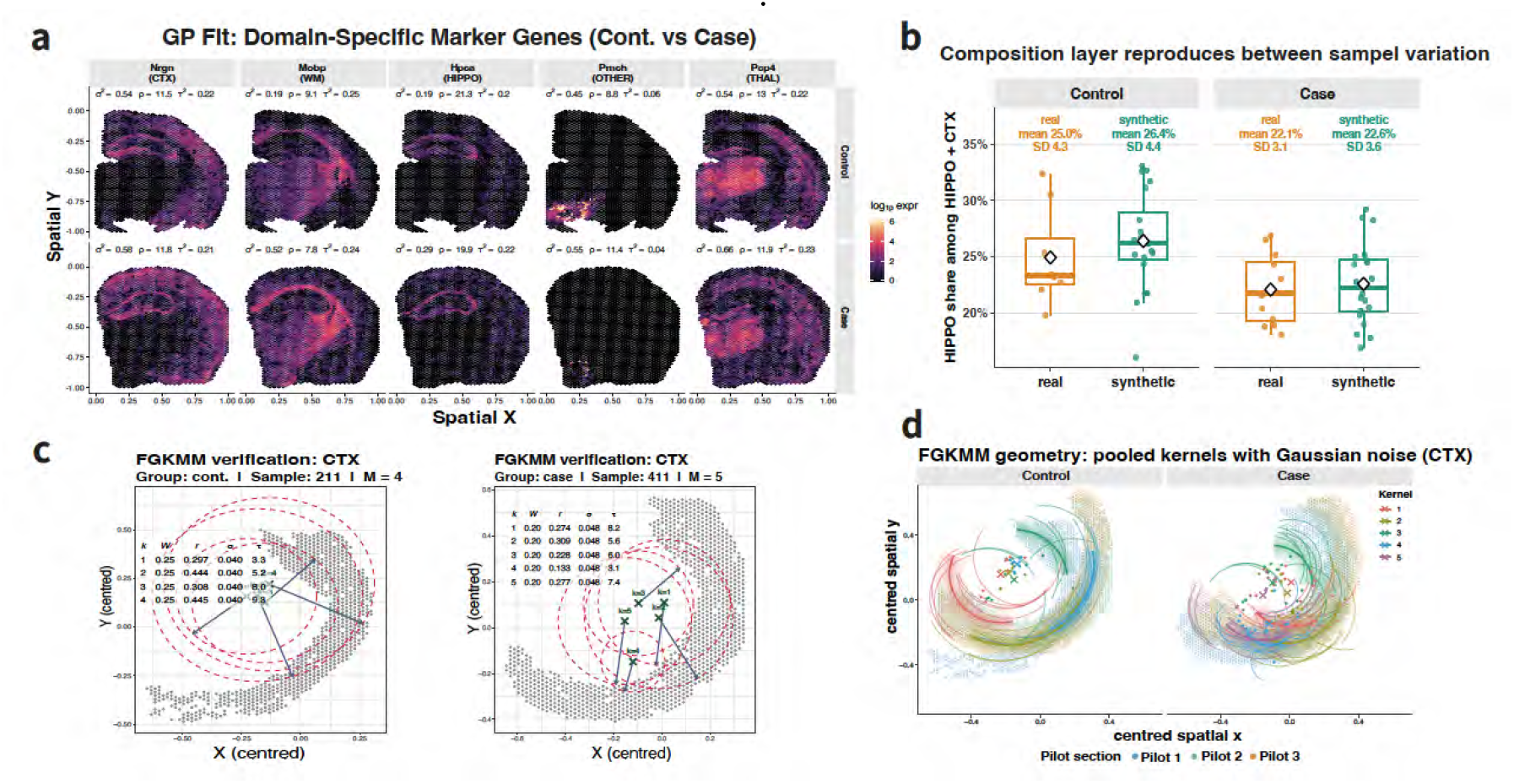
The three layers fit a second cohort and its between-sample variation (5xFAD Visium, five domains, (*T, R*) = hippocampus (HIPPO) versus cortex (CTX)). **(a)** Expression layer 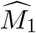. Spatial log-expression of five domain markers, *Nrgn* (CTX), *Mobp* (white matter), *Hpca* (HIPPO), *Pmch* (other), and *Pcp4* (thalamus), in control (top) and case (bottom), with the fitted Gaussian-process partial sill *σ*^2^, range *ρ*, and nugget *τ* ^2^ per group. **(b)** Composition layer 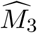. The HIPPO share within the (*T, R*) pair matches the real cohort in the synthetic cohort in mean and between-sample spread, control 25.0% against 26.4% at s.d. 4.3 against 4.4, and case 22.1% against 22.6% at s.d. 3.1 against 3.6 (percentage points; diamonds, group means). **(c)** Geometry layer 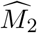, single samples. FGKMM fit of the CTX domain in one control sample (211, *M* = 4 kernels) and one case sample (411, *M* = 5), with per-kernel weight *W*, radius *r*, radial dispersion *σ*, and angular concentration *τ* . These kernel parameters are local to the geometry layer and are distinct from the Gaussian-process parameters in **a. (d)** Geometry layer 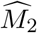, pooled. FGKMM kernels pooled across the three pilot samples capture the between-sample variation in CTX position, orientation, and spread, for control and case.

**Extended Data Figure 2.**
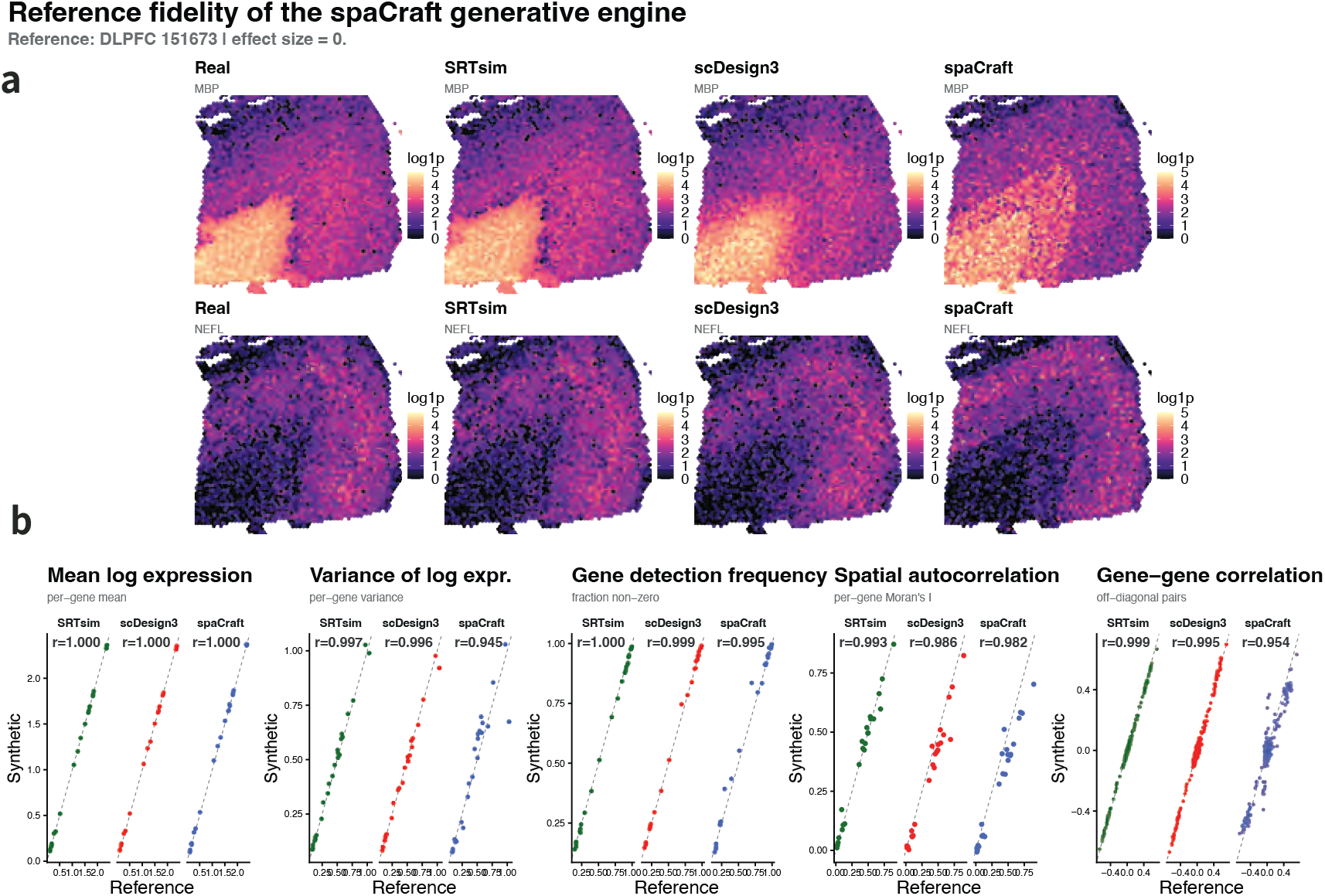
Pilot fidelity of the generator on the DLPFC reference sample. **(a)** Synthetic expression maps for *MBP* and *NEFL* against the real tissue and two single-sample simulators (SRTsim, scDesign3). **(b)** Per-gene scatter of five summaries, the mean, variance, detection frequency, Moran’s *I*, and gene–gene correlation, against the reference, with Pearson *r* of 1.00, 0.94, 0.99, 0.98, and 0.95. The spaCraft generator attains fidelity comparable to the two dedicated single-sample simulators.

**Extended Data Figure 3.**
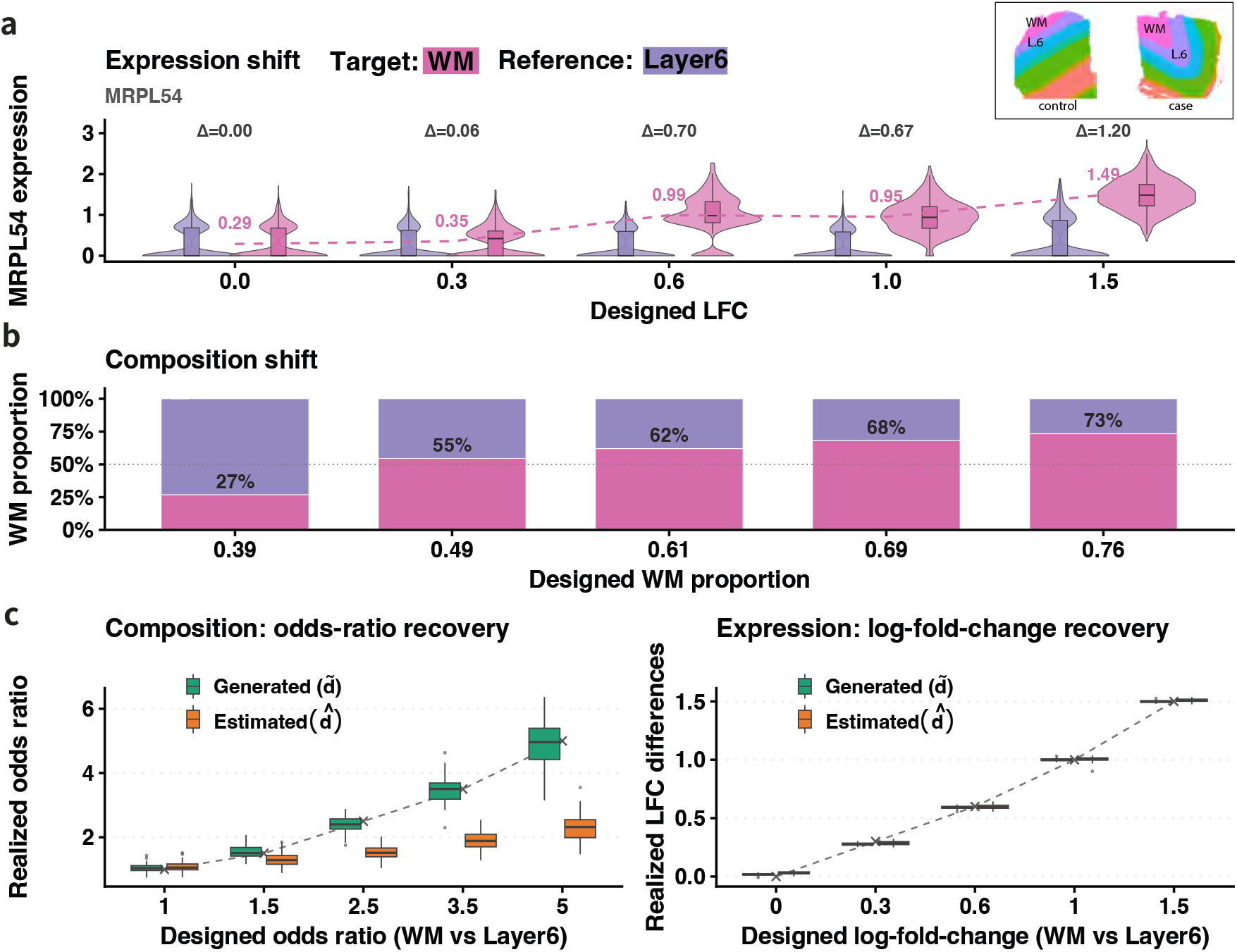
Effect realization under the generator labels and analysis-aware effect fidelity under the recovered labels. **(a)** Expression shift of a marker gene (*MRPL54* ) between white matter (WM) and Layer 6 across specified log-fold-changes Δ. **(b)** Compositional shift, the realized WM share within the (WM, Layer 6) pair across specified target proportions. **(c)** Analysis-aware effect fidelity on both scales, comparing the value realized under the generator labels 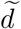 (green) with that under the recovered labels 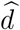 (orange). The odds ratio (left) is attenuated under 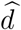, the loss expected from clustering error, whereas the log-fold-change (right) tracks its specified value under both.

**Extended Data Figure 4.**
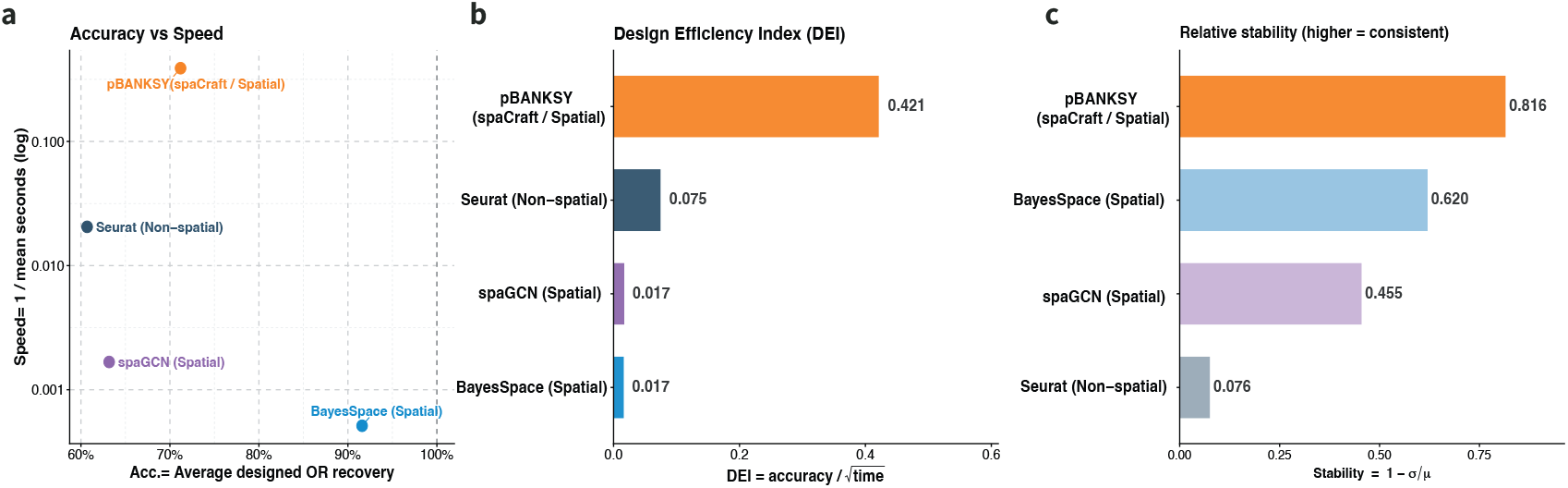
Benchmark of candidate recovery procedures within spaCraft. pBANKSY, BayesSpace, SpaGCN, and Seurat were each substituted into the same recovery step of the spaCraft loop on synthetic DLPFC cohorts. **(a)** Accuracy of recovering the specified effect against speed (inverse mean runtime, log scale). **(b)** Design efficiency index, accuracy per unit runtime, each value as a percentage of the best. **(c)** Run-to-run stability of the recovered partition, higher being more consistent. These results motivated the use of pBANKSY as the default recovery module in spaCraft.

**Extended Data Figure 5.**
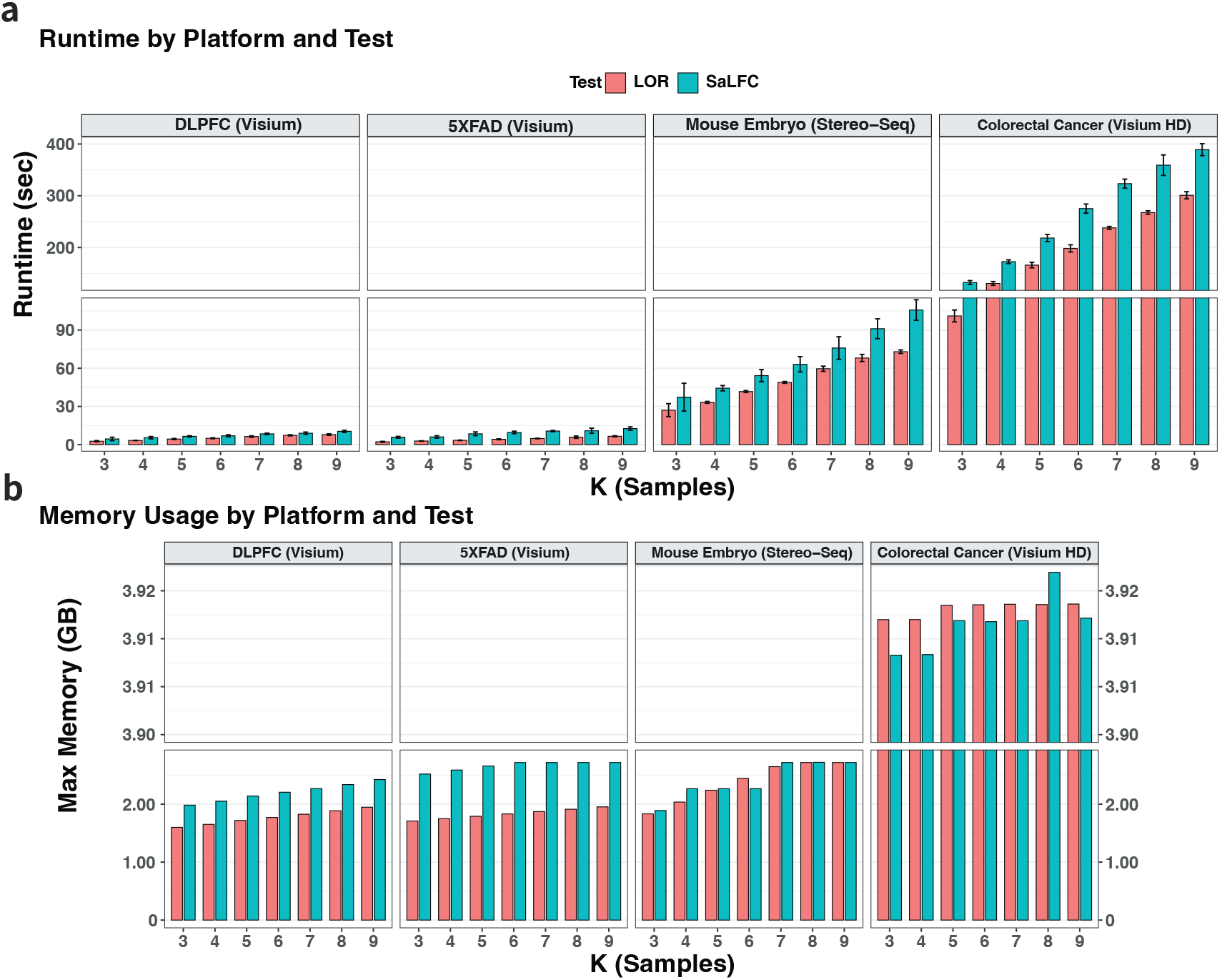
Runtime and memory of LOR and SaLFC across platforms. **(a)** Runtime (seconds) on four datasets, DLPFC (Visium), 5xFAD (Visium), mouse embryo (Stereo-seq), and colorectal cancer (Visium HD), against sample size *K*, with bars the mean and error bars the mean ± s.d. across replicates. **(b)** Maximum memory (MB) for the same evaluations. Broken *y*-axes accommodate the larger footprint of the Visium HD dataset. All measurements were made on the Ohio Supercomputer Center Ascend cluster using 40 CPU cores.

**Extended Data Figure 6.**
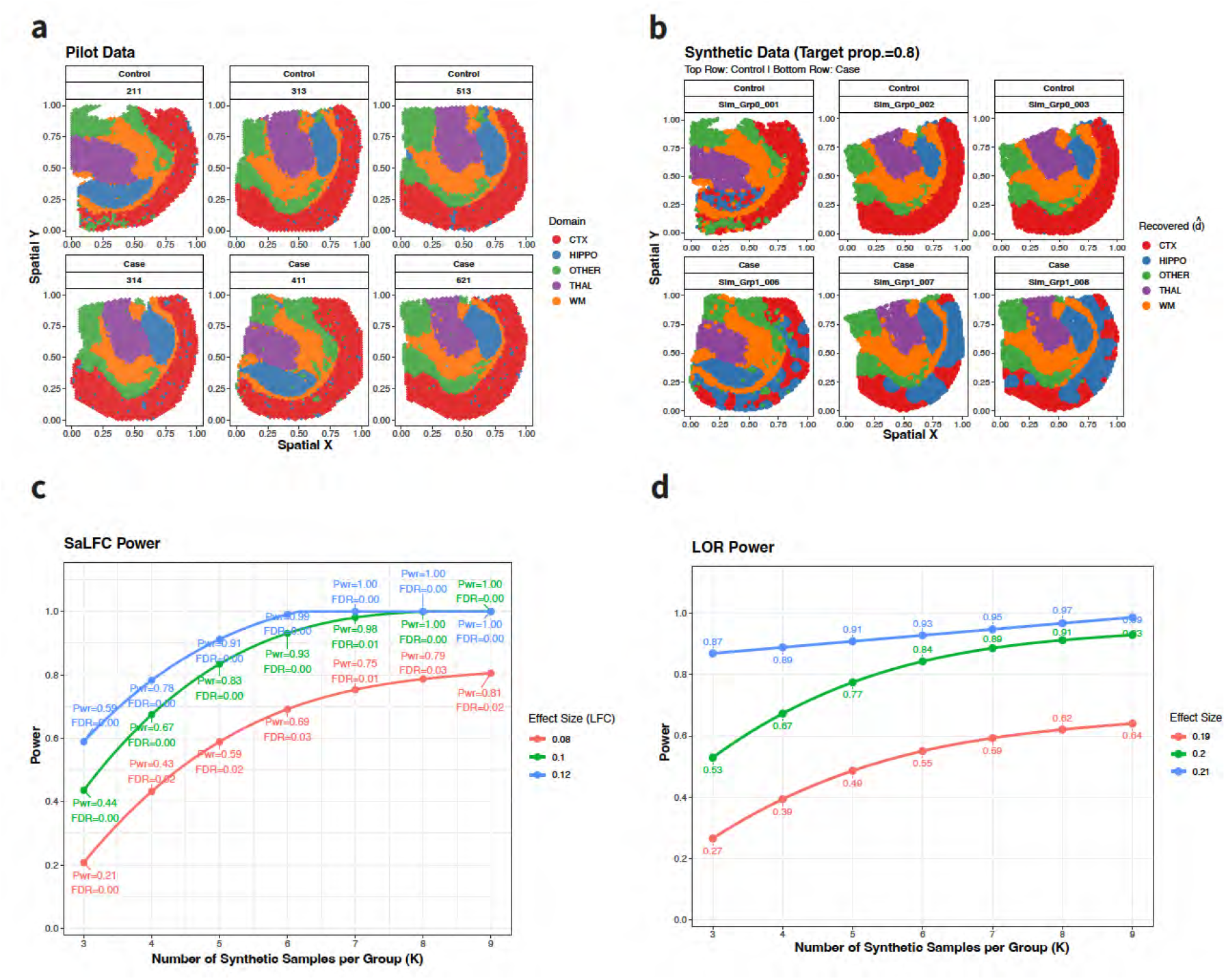
In-loop domain recovery yields calibrated, endpoint-specific power on a second cohort (5xFAD Visium, (*T, R*) = hippocampus (HIPPO) versus cortex (CTX)). **(a)** Annotated domains of the six pilot samples, three control (211, 313, 513) and three case (314, 411, 621), across five domains (CTX, HIPPO, thalamus (THAL), white matter (WM), and other). **(b)** Domains recovered by pBANKSY on synthetic samples generated at a case target share of 0.8, after optimal-transport rearrangement, for control (top) and case (bottom). **(c)** SaLFC power against the per-group sample size *K* for three injected log-fold-changes (0.08, 0.10, 0.12), with power and empirical FDR annotated at each point. **(d)** LOR power against *K* for three case target shares (0.19, 0.20, 0.21). The effect grids differ from those of the DLPFC analysis because the two cohorts differ in baseline target–reference separation and in baseline target share.

**Extended Data Table 1.**
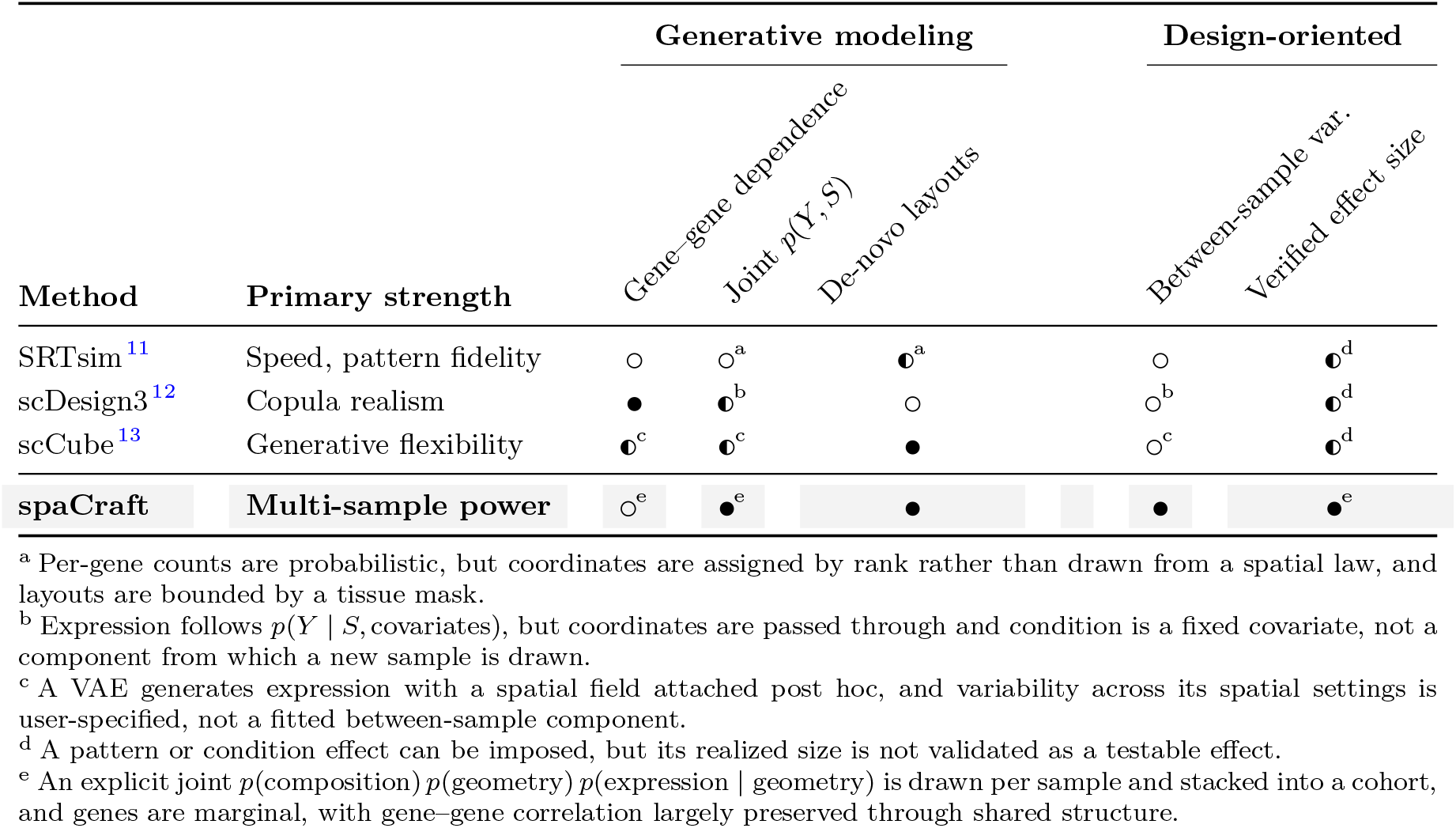
spaCraft as a generator, alongside dedicated spatial simulators. Each simulator excels at reproducing an observed sample, SRTsim for speed and spatial-pattern fidelity, scDesign3 for a copula over genes, and scCube for deep-generative flexibility. spaCraft is built for a different task, generating a multi-sample cohort for a power analysis. This column records whether gene–gene dependence is modeled explicitly, which neither SRTsim nor spaCraft does, although resampling and shared structure respectively preserve much of the observed correlation. spaCraft is the only fully probabilistic joint generator of expression and coordinates that carries a between-sample variance component and realizes a user-specified effect at a verified size. Symbols: addressed, partial, not addressed.

**Extended Data Table 2.** Overview of the spatial transcriptomics datasets. The four datasets used for pilot-driven power analysis in spaCraft. Columns give the platform, the number of samples in the control (*K*_0_) and case (*K*_1_) groups of the full cohort, the range of spots or bins per sample, the number of genes retained after quality control, the number of annotated domains, the target and reference domains analyzed, and the inmemory object size in R. Pilots used within the design loop are subsets of these cohorts, three samples per group unless stated otherwise.

| Dataset | Platform | $K_0 / K_1$ | Spots per sample | Genes | Domains | Target vs. reference | Object size |
| --- | --- | --- | --- | --- | --- | --- | --- |
| DLPFC | Visium | 6 / 6 | 3,431–4,788 | 33,538 | 7 | WM vs. Layer 6 | 2.0 GB |
| 5xFAD | Visium | 8 / 12 | 2,206–3,163 | 32,285 | 5 | Hippocampus vs. cortex | 3.4 GB |
| Mouse embryo | Stereo-seq | 3 / 3 | 5,457–14,991 | 14,611–15,954 | 5 | Lung vs. neural | 9.0 GB |
| Colorectal cancer | Visium HD | 2 / 3 | 62,161–127,471 | 12,106–12,866 | 6 | Normal epithelium vs. stroma | 12.2 GB |

