## Supplementary note for "spaCraft: calibrated power analysis and sample-size planning for multi-sample spatial transcriptomics"

August 28, 2026

### S1 Gene-set construction

Three pilot-derived gene sets are fixed before simulation and held constant across Monte Carlo replicates. Throughout this section, sample indices are suppressed when they are not required. We use $y_{ig} = \log(1 + X_{ig})$, $\mathcal{I}_d = \{i : d_i = d\}$, and $\mathcal{I}_{\setminus d} = \{i : d_i \neq d\}$.

#### S1.1 Labeling set $\mathcal{G}_{\text{SVG}}$

Domain-informative genes are identified from user-selected representative pilot samples. For domain $d$, let $m_{g,\setminus d}$ be the median of $\{y_{ig} : i \in \mathcal{I}_{\setminus d}\}$ and define $\mathcal{I}^{\text{low}}_{\setminus d}(g) = \left\{i \in \mathcal{I}_{\setminus d} : y_{ig} \leq m_{g,\setminus d}\right\}$. We compare the within-domain mean with the lower-truncated off-domain mean,

$$\Delta^{\text{low}}_{g,d} = \bar{y}_{g,d} - \bar{y}^{\text{low}}_{g,\setminus d}, \qquad \bar{y}_{g,d} = \frac{1}{|\mathcal{I}_d|}\sum_{i\in\mathcal{I}_d} y_{ig}, \qquad \bar{y}^{\text{low}}_{g,\setminus d} = \frac{\sum_{i\in\mathcal{I}^{\text{low}}_{\setminus d}(g)} y_{ig}}{|\mathcal{I}^{\text{low}}_{\setminus d}(g)|}. \tag{S1}$$

For each domain, $\mathcal{S}_d = \left\{g : \bar{y}_{g,d} \geq \tau_{\text{mean}}, \quad |\Delta^{\text{low}}_{g,d}| \geq \tau_{\text{LFC}}\right\}$. Genes are ranked by $|\Delta^{\text{low}}_{g,d}|$, and the top $M$ per domain are retained and combined across domains and representative samples. The thresholds $\tau_{\text{mean}}$ and $\tau_{\text{LFC}}$, and the truncation level $M$, are user-specified and may be chosen data-adaptively.

Let $\mathcal{G}^{\text{base}}_{\text{SVG}}$ denote the resulting domain-informative marker set. The final expression set supplied to synthetic generation and domain recovery is

$$\mathcal{G}_{\text{SVG}} = \mathcal{G}^{\text{base}}_{\text{SVG}} \cup \mathcal{G}_{\text{Null}}, \tag{S2}$$

where $\mathcal{G}_{\text{Null}}$ is defined below.

#### S1.2 Empirical-null set $\mathcal{G}_{\text{Null}}$

The empirical-null set contains genes that are sufficiently expressed but spatially stable in the control-group pilot samples. For sample $(k, 0)$, let $\bar{y}^{(k,0)}_{g,\cdot} = \frac{1}{N^{(k,0)}}\sum_i y^{(k,0)}_{ig}$. The principal filters require

$$\bar{y}_{g,\cdot}^{(k,0)} \geq \tau_{\text{global}}, \qquad \max_{d \in \mathcal{A}} |\Delta_{g,d}^{\text{low},(k,0)}| \leq \tau_{\text{stable}}. \tag{S3}$$

The implementation additionally requires adequate detection across domains and low variation among the domain-specific means. Genes must satisfy the spatial-stability criteria in every available control pilot sample. By default, genes showing a control–case difference in global mean expression greater than a user-specified tolerance are also excluded.

The remaining genes are ranked by between-group stability, followed by across-domain homogeneity and residual spatial contrast, and the top $M_{\text{null}}$ define $\mathcal{G}_{\text{Null}}$. All filtering thresholds and $M_{\text{null}}$ are user-specified and may be chosen data-adaptively for each dataset.

##### S1.3 Injection set $\mathcal{G}_{\text{spike}}$

For the prespecified target and reference domains $(T, R)$, the selected pilot sample defines

$$\Delta_g^{TR} = \bar{y}_{g,T} - \bar{y}_{g,R}, \qquad g \in \mathcal{G}_{\text{Null}}. \tag{S4}$$

The injection set consists of the $n_{\text{de}}$ genes in $\mathcal{G}_{\text{Null}}$ with the largest $|\Delta_g^{TR}|$, $\mathcal{G}_{\text{spike}} \subset \mathcal{G}_{\text{Null}}$. Thus, injected and noninjected genes originate from the same pilot-defined stable population and differ only by the imposed expression shift. The set size $n_{\text{de}}$ is user-specified and may be chosen data-adaptively for each dataset.

##### S1.4 Conservativeness of the injection-set selection

Selecting $\mathcal{G}_{\text{spike}}$ from stable genes with the largest baseline $|\Delta_g^{TR}|$ deliberately avoids an overly favorable setting in which the target and reference domains are nearly indistinguishable before injection. Such genes are also more sensitive to contamination caused by imperfect domain recovery, which attenuates their target–reference contrast and adds between-sample variability. The resulting power evaluation is therefore stringent with respect to recovery error and tends to favor larger, rather than smaller, sample-size recommendations.

#### S2 Spatial gene expression model

This note expands the expression model $M_1$ described in Methods. It specifies the joint variation of the target and reference domain means, the per-sample fitting and within-group pooling, and the interpretation of the fitted spatial parameters.

##### S2.1 Additive decomposition

For gene $g$ at spot $i$ of sample $(k, c)$, log expression is decomposed as

$$y_{ig}^{(k,c)} = \mu_{g,d_i}^{(k,c)} + B_g^{(k,c)} + \eta_{ig}^{(k,c)} + \epsilon_{ig}^{(k,c)}, \tag{S5}$$

where $\mu_{g,d}^{(k,c)}$ is the sample- and domain-specific mean, $B_g^{(k,c)} \sim \mathcal{N}(0, \sigma_{B,g,c}^2)$ is a sample-level shift shared across domains, $\boldsymbol{\eta}_g^{(k,c)}$ is a mean-zero GP, and $\epsilon_{ig}^{(k,c)} \sim \mathcal{N}(0, \tau_{g,c}^2)$ is independent spot-level noise. Conditional on the domain means and labels, the three random components are mutually independent.

The GP has isotropic exponential covariance

$$\boldsymbol{\eta}_g^{(k,c)} \sim \mathcal{N}\left(\mathbf{0}, \boldsymbol{\Sigma}_g^{(k,c)}\right), \qquad \boldsymbol{\Sigma}_g^{(k,c)}(i,j) = \sigma_{g,c}^2 \exp\left(-\frac{\|\mathbf{s}_i - \mathbf{s}_j\|_2}{\rho_{g,c}}\right). \tag{S6}$$

Thus, $\operatorname{Cov}\left(y_{ig}^{(k,c)}, y_{jg}^{(k,c)} \mid \boldsymbol{\mu}_g^{(k,c)}, \mathbf{d}\right) = \boldsymbol{\Sigma}_g^{(k,c)}(i,j) + \tau_{g,c}^2 \mathbb{1}\{i = j\}$, where $\sigma_{g,c}^2$ is the partial sill, $\rho_{g,c}$ the range, and $\tau_{g,c}^2$ the nugget variance. These parameters are group-specific and estimated separately within each group.

For the prespecified target and reference domains $(T, R)$, the sample-specific domain means are modeled jointly as

$$\begin{pmatrix} \mu_{g,T}^{(k,c)} \\ \mu_{g,R}^{(k,c)} \end{pmatrix} \sim \mathcal{N}\left(\begin{pmatrix} \mu_{g,T,c} \\ \mu_{g,R,c} \end{pmatrix}, \sigma_{\text{bio},g,c}^2 \begin{pmatrix} 1 & \varrho_{TR,g,c} \\ \varrho_{TR,g,c} & 1 \end{pmatrix}\right). \tag{S7}$$

Their between-sample contrast variance is therefore

$$\operatorname{Var}\left(\mu_{g,T}^{(k,c)} - \mu_{g,R}^{(k,c)}\right) = 2\sigma_{\text{bio},g,c}^2 \left(1 - \varrho_{TR,g,c}\right). \tag{S8}$$

Because $B_g^{(k,c)}$ is shared across domains, it cancels from the target–reference contrast. A positive $\varrho_{TR,g,c}$ captures co-movement of the two domain means across samples and reduces their contrast variance.

#### S2.2 Per-sample fitting and within-group pooling

The model is fitted to each pilot sample and then pooled within group. For domain $d$ in sample $(k, c)$, let

$$\bar{y}_{g,d}^{(k,c)} = \frac{1}{n_d^{(k,c)}} \sum_{i:d_i=d} y_{ig}^{(k,c)}, \qquad \widehat{\mu}_{g,d,c} = \frac{1}{K_c} \sum_{k=1}^{K_c} \bar{y}_{g,d}^{(k,c)}, \tag{S9}$$

where $n_d^{(k,c)}$ is the number of spots assigned to domain $d$ in sample $(k, c)$.

The sample-level shift is estimated from the average deviation of the sample-specific domain means from their group levels,

$$\widehat{B}_g^{(k,c)} = \frac{1}{D} \sum_{d \in \mathcal{A}} \left(\bar{y}_{g,d}^{(k,c)} - \widehat{\mu}_{g,d,c}\right), \qquad \widehat{\sigma}_{B,g,c}^2 = \operatorname{Var}_k\left(\widehat{B}_g^{(k,c)}\right). \tag{S10}$$

The corresponding shift-adjusted domain mean is $\widehat{\mu}_{g,d}^{(k,c)} = \bar{y}_{g,d}^{(k,c)} - \widehat{B}_g^{(k,c)}$. The biological variance and target–reference correlation are estimated by

$$\widehat{\sigma}_{\text{bio},g,c}^2 = \frac{1}{D} \sum_{d \in \mathcal{A}} \operatorname{Var}_k\left(\widehat{\mu}_{g,d}^{(k,c)}\right), \qquad \widehat{\varrho}_{TR,g,c} = \operatorname{Cor}_k\left(\widehat{\mu}_{g,T}^{(k,c)}, \widehat{\mu}_{g,R}^{(k,c)}\right), \tag{S11}$$

where $\operatorname{Var}_k$ and $\operatorname{Cor}_k$ denote empirical variance and correlation over $k = 1, \ldots, K_c$. Removing $\widehat{B}_g^{(k,c)}$ prevents the global sample shift from being counted again as domain-specific biological variation.

For spatial covariance fitting, each pilot sample is centered by its sample-specific domain means. For each gene, the centered expression $y_{ig}^{(k,c)} - \bar{y}_{g,d_i}^{(k,c)}$ is fitted by maximum likelihood under the nearest-neighbor GP approximation implemented in BRISC[1,2]. Let

$$\widehat{\boldsymbol{\theta}}_g^{(k,c)} = \left(\widehat{\sigma}_g^{2,(k,c)}, \widehat{\rho}_g^{(k,c)}, \widehat{\tau}_g^{2,(k,c)}\right)$$

denote the resulting sample-specific covariance estimates. The generative parameters are their within-group averages,

$$\widehat{\boldsymbol{\theta}}_{g,c} = \frac{1}{K_c}\sum_{k=1}^{K_c}\widehat{\boldsymbol{\theta}}_g^{(k,c)} = \left(\widehat{\sigma}_{g,c}^2, \widehat{\rho}_{g,c}, \widehat{\tau}_{g,c}^2\right). \tag{S12}$$

An intercept is retained in each spatial fit to absorb residual centering error. The empirical quantities $\widehat{\sigma}^2_{B,g,c}$, $\widehat{\sigma}^2_{\mathrm{bio},g,c}$, and $\widehat{\varrho}_{TR,g,c}$ require replicated samples and are estimated when $K_c \geq 2$.

##### S2.3 Identifiability and cross-sample transfer

Cross-sample transfer was strongest for the partial sill, nugget variance, and the microergodic ratio $\sigma^2_{g,c}/\rho_{g,c}$, whereas the range $\rho_{g,c}$ transferred less reliably on its own (Fig. 2f). This agrees with fixed-domain asymptotics, under which $\sigma^2$ and $\rho$ are not separately consistently estimable for an exponential covariance, while $\sigma^2/\rho$ is identifiable[3]. We therefore retain $(\widehat{\sigma}^2_{g,c}, \widehat{\rho}_{g,c})$ as the working covariance parameterization, but assess cross-sample stability through $\widehat{\sigma}^2_{g,c}/\widehat{\rho}_{g,c}$. During generation, $\widehat{\rho}_{g,c}$ controls spatial decay and the spatial field is rescaled to marginal variance $\widehat{\sigma}^2_{g,c}$ before adding nugget noise (Supplementary Note S5).

#### S3 Spatial geometry model

The geometry model $M_2$ separates between-sample variation in domain placement from variation in within-domain shape. Pilot coordinates are normalized to $[0,1]^2$ within each sample. For domain $d$ of sample $(k,c)$, define

$$\widehat{\mathbf{G}}_d^{(k,c)} = \frac{1}{n_d^{(k,c)}}\sum_{i:d_i=d}\mathbf{s}_i^{(k,c)}, \qquad \mathbf{s}_i'^{(k,c)} = \mathbf{s}_i^{(k,c)} - \widehat{\mathbf{G}}_d^{(k,c)}. \tag{S13}$$

##### S3.1 Domain placement

Pilot centroids are modeled within each group as

$$\widehat{\mathbf{G}}_d^{(k,c)} \sim \mathcal{N}\left(\boldsymbol{\mu}_{G,d}^{(c)}, \boldsymbol{\Sigma}_{G,d}^{(c)}\right), \tag{S14}$$

with

$$\widehat{\boldsymbol{\mu}}_{G,d}^{(c)} = \frac{1}{K_c}\sum_{k=1}^{K_c}\widehat{\mathbf{G}}_d^{(k,c)}, \qquad \widehat{\boldsymbol{\Sigma}}_{G,d}^{(c)} = \mathrm{Cov}_k\left(\widehat{\mathbf{G}}_d^{(k,c)}\right), \tag{S15}$$

for $K_c \geq 2$. Thus, $\widehat{\boldsymbol{\Sigma}}_{G,d}^{(c)}$ captures between-sample variation in domain position.

##### S3.2 Within-domain Fisher–Gaussian kernel mixture

Conditional on its centroid, the centered coordinates of domain $d$ are modeled by an FGKMM[4],

$$f_d^{(k,c)}(\mathbf{s}') = \sum_{m=1}^{M_d} \omega_{m,d}^{(k,c)} \mathrm{FG}_{\sigma_d^{(k,c)}} \left( \mathbf{s}' \mid \boldsymbol{\theta}_{m,d}^{(k,c)} \right), \qquad \sum_{m=1}^{M_d} \omega_{m,d}^{(k,c)} = 1, \tag{S16}$$

where $\boldsymbol{\theta}_{m,d}^{(k,c)} = (\mathbf{c}, r, \boldsymbol{\phi}, \tau)$. Here, $\mathbf{c} \in \mathbb{R}^2$ and $r > 0$ define a local center and radius, $\boldsymbol{\phi} \in \mathbb{S}^1$ the direction, $\tau \geq 0$ the angular concentration, and $\sigma > 0$ the radial dispersion. The component density is

$$\mathrm{FG}_\sigma \left( \mathbf{s}' \mid \boldsymbol{\theta} \right) = \frac{C_2(\tau)}{C_2 \left( \| \tau \boldsymbol{\phi} + r \mathbf{s}' / \sigma^2 \| \right)} (2\pi\sigma^2)^{-1} \exp \left[ - \frac{\| \mathbf{s}' - \mathbf{c} \|_2^2 + r^2}{2\sigma^2} \right], \tag{S17}$$

where $C_2(\tau) = \{2\pi I_0(\tau)\}^{-1}$[5]. Mixtures of these kernels accommodate irregular, non-convex, and disconnected domains. The geometry parameters $\sigma$ and $\tau$ are distinct from their expression-model counterparts in Supplementary Note S2.

##### S3.3 Model selection and within-group pooling

For each group-domain pair, the component count is selected from the pooled centered pilot coordinates by

$$\widehat{M}_d = \underset{M \in \{3, \ldots, 6\}}{\operatorname{argmin}} \left\{ -2\widehat{\ell}_d(M) + 6M \log N_d \right\}, \tag{S18}$$

where $N_d$ is the number of pooled spots. Each pilot sample is then refitted at the common order $\widehat{M}_d$, allowing its fitted kernel parameters to vary across samples under a fixed model dimension. The sample-specific parameters are pooled within group using

$$\begin{aligned}
\mathbf{c}_{m,d}^{(k,c)} &\sim \mathcal{N}_2 \left( \boldsymbol{\mu}_{c,m,d}^{(c)}, \boldsymbol{\Sigma}_{c,m,d}^{(c)} \right), & r_{m,d}^{(k,c)} &\sim \mathcal{N} \left( \mu_{r,m,d}^{(c)}, v_{r,m,d}^{(c)} \right) \mathbb{1}\{r > 0\}, \\
\boldsymbol{\phi}_{m,d}^{(k,c)} &\sim \mathrm{vMF}_{\mathbb{S}^1} \left( \boldsymbol{\mu}_{\phi,m,d}^{(c)}, \kappa_{\phi,m,d}^{(c)} \right), & \tau_{m,d}^{(k,c)} &\sim \mathrm{Gamma} \left( a_{\tau,m,d}^{(c)}, b_{\tau,m,d}^{(c)} \right), \\
\sigma_d^{2(k,c)} &\sim \text{Inv-Gamma} \left( a_{\sigma,d}^{(c)}, b_{\sigma,d}^{(c)} \right), & \boldsymbol{\omega}_d^{(k,c)} &\sim \mathrm{Dirichlet} \left( \boldsymbol{\alpha}_d^{(c)} \right).
\end{aligned} \tag{S19}$$

The hyperparameters are estimated from the corresponding per-sample fits, thereby propagating pilot variation in position, orientation, spread, and mixture weights into synthetic samples.

##### S3.4 Generation from the pooled geometry model

For synthetic sample $(k, c)$, domain $d$ first receives a centroid

$$\widetilde{\mathbf{G}}_d^{(k,c)} \sim \mathcal{N} \left( \widehat{\boldsymbol{\mu}}_{G,d}^{(c)}, \widehat{\boldsymbol{\Sigma}}_{G,d}^{(c)} \right). \tag{S20}$$

Kernel weights and parameters are then drawn from the fitted group-level distributions in Eq. (S19). To preserve pilot geometry, component centers and directions use a cloudanchored draw,

$$\widetilde{\mathbf{c}} = \lambda_{\text{cloud}}\mathbf{c}_j + (1-\lambda_{\text{cloud}})\mathbf{c}_{\text{pool}}, \qquad \widetilde{\boldsymbol{\phi}} = \frac{\lambda_{\text{cloud}}\boldsymbol{\phi}_j + (1-\lambda_{\text{cloud}})\boldsymbol{\phi}_{\text{pool}}}{\left\|\lambda_{\text{cloud}}\boldsymbol{\phi}_j + (1-\lambda_{\text{cloud}})\boldsymbol{\phi}_{\text{pool}}\right\|_2}, \tag{S21}$$

where $j$ indexes a pilot fit and $\lambda_{\text{cloud}} \in [0,1]$ controls the degree of anchoring. Given a sampled component,

$$\mathbf{u} \sim \text{vMF}_{\mathbb{S}^1}(\widetilde{\boldsymbol{\phi}}, \widetilde{\tau}), \qquad \widetilde{\mathbf{s}}'_i = \widetilde{\mathbf{c}} + \widetilde{r}\mathbf{u} + \boldsymbol{\varepsilon}_i, \qquad \boldsymbol{\varepsilon}_i \sim \mathcal{N}(\mathbf{0}, \widetilde{\sigma}^2\mathbf{I}_2), \tag{S22}$$

and the absolute synthetic coordinate is $\widetilde{\mathbf{s}}_i^{(k,c)} = \widetilde{\mathbf{G}}_d^{(k,c)} + \widetilde{\mathbf{s}}'_i$. Repeating this draw for the domain count supplied by $\widehat{M}_3$ generates the synthetic point cloud and its generator labels $\widetilde{d}_i$.

### S4 Domain composition model

The composition model $M_3$ describes the abundance of the target domain relative to the prespecified reference domain while preserving the remaining tissue composition.

#### S4.1 Reference-anchored subcomposition

For sample $(k,c)$, let

$$m^{(k,c)} = n_T^{(k,c)} + n_R^{(k,c)}, \qquad N^{(k,c)} = \sum_{d\in\mathcal{A}} n_d^{(k,c)}$$

. If $\pi_d^{(k,c)}$ denotes the underlying abundance of domain $d$, the target share within the $(T,R)$ subcomposition is

$$p^{(k,c)} = \frac{\pi_T^{(k,c)}}{\pi_T^{(k,c)} + \pi_R^{(k,c)}}, \qquad \text{logit}\left(p^{(k,c)}\right) = \log\frac{\pi_T^{(k,c)}}{\pi_R^{(k,c)}}. \tag{S23}$$

Conditional on $m^{(k,c)}$,

$$n_T^{(k,c)} \sim \text{Binomial}\left(m^{(k,c)}, p^{(k,c)}\right), \qquad n_R^{(k,c)} = m^{(k,c)} - n_T^{(k,c)}. \tag{S24}$$

Thus, the target–reference contrast is invariant to the abundances of the remaining domains. Let $\mathcal{A}_{-TR} = \mathcal{A}\setminus\{T,R\}$. The residual spot budget is allocated by

$$\left(n_d^{(k,c)}\right)_{d\in\mathcal{A}_{-TR}} \sim \text{Multinomial}\left(N^{(k,c)} - m^{(k,c)}, \boldsymbol{\pi}_{-TR}^{(c)}\right). \tag{S25}$$

#### S4.2 Pilot fitting

The group-level marginal target shares are modeled by

$$\text{logit}(p_c) = \beta_0 + \beta_1 c, \qquad c \in \{0,1\}, \tag{S26}$$

and estimated jointly from the pilot pairs $\{(n_T^{(k,c)}, m^{(k,c)})\}$ as described in Methods equation (8). The resulting $\widehat{p}_0$ and $\widehat{p}_1$ are the fitted control and case target shares, and $\exp(\widehat{\beta}_1)$

is the corresponding pilot odds ratio. For $d \in \mathcal{A}_{-TR}$, the background probabilities are estimated from the average residual-domain proportions,

$$\widehat{\pi}_d^{(c)} = \frac{q_d^{(c)}}{\sum_{d' \in \mathcal{A}_{-TR}} q_{d'}^{(c)}}, \qquad q_d^{(c)} = \frac{1}{K_c} \sum_{k=1}^{K_c} \frac{n_d^{(k,c)}}{N^{(k,c)} - m^{(k,c)}}. \tag{S27}$$

The pair-budget moments are

$$\widehat{\mu}_{m,c} = \frac{1}{K_c} \sum_{k=1}^{K_c} m^{(k,c)}, \qquad \widehat{\sigma}_{m,c}^2 = \mathrm{Var}_k \left( m^{(k,c)} \right), \tag{S28}$$

and are preserved during synthetic generation.

##### S4.3 Between-sample variation in target abundance

To reproduce between-sample heterogeneity beyond Binomial sampling, synthetic target shares follow

$$\widetilde{p}^{(k,c)} = \mathrm{logit}^{-1} \left( a_c + u^{(k,c)} \right), \qquad u^{(k,c)} \sim \mathcal{N}\left(0, \sigma_{\mathrm{re}}^2\right). \tag{S29}$$

Because the logistic transformation changes the marginal mean, $a_c$ is calibrated numerically so that

$$\mathrm{E}\left[\widetilde{p}^{(k,c)}\right] = p_c. \tag{S30}$$

The common scale $\sigma_{\mathrm{re}}$ is calibrated from the observed between-sample variation of the pilot logit shares,

$$\widehat{\sigma}_{\mathrm{re}} = \frac{1}{2} \sum_{c \in \{0,1\}} \mathrm{sd}_k \left[ \mathrm{logit} \left( \frac{n_T^{(k,c)}}{m^{(k,c)}} \right) \right], \tag{S31}$$

with a small numerical truncation applied when a pilot share is 0 or 1. This preserves the marginal group share while propagating the observed between-sample compositional spread.

#### S5 Synthetic data generation

A synthetic sample $\widetilde{\mathcal{D}}^{(k,c)} = (\widetilde{\mathbf{X}}^{(k,c)}, \widetilde{\mathbf{S}}^{(k,c)}, \widetilde{\mathbf{d}}^{(k,c)})$ is generated in the order $\widehat{M}_3 \to \widehat{M}_2 \to \widehat{M}_1$. The composition model determines domain counts, the geometry model generates continuous coordinates and generator labels, and the expression model generates spatially structured expression conditional on these labels.

##### S5.1 Pilot-conditioned spatial texture

To retain local spatial features not captured by the parametric GP, a pilot sample is drawn from the same group. For a synthetic spot $i$ with $\widetilde{d}_i = d$, let $\mathcal{N}_d(i)$ denote its $k_{\mathrm{cond}}$ nearest pilot spots annotated as domain $d$. The pilot-conditioned field is

$$\widetilde{\eta}_{ig}^{\mathrm{cond}} = \sum_{j \in \mathcal{N}_d(i)} \bar{w}_{ij} \left( y_{jg}^{\mathrm{pilot}} - \bar{y}_{g,d}^{\mathrm{pilot}} \right), \qquad \bar{w}_{ij} = \frac{\exp(-\|\widetilde{\mathbf{s}}_i - \mathbf{s}_j^{\mathrm{pilot}}\|_2^2 / h_d)}{\sum_{j' \in \mathcal{N}_d(i)} \exp(-\|\widetilde{\mathbf{s}}_i - \mathbf{s}_{j'}^{\mathrm{pilot}}\|_2^2 / h_d)}. \tag{S32}$$

The bandwidth $h_d$ is determined by the local neighbor distances, and the resulting field is centered gene-wise so that it contributes spatial texture without changing the generated domain means.

##### S5.2 Scalable parametric spatial field

Rather than drawing an exact $N$-dimensional GP for every gene, spaCraft uses a scalable graph- or basis-based approximation. In graph mode (used for very high resolution platform (e.g., Visium-HD),

$$\widetilde{\eta}_{ig}^{\text{para}} = \sum_{j \in \mathcal{N}(i) \cup \{i\}} \bar{w}_{ij} z_j, \qquad w_{ij} = \exp\left(-\frac{\|\widetilde{\mathbf{s}}_i - \widetilde{\mathbf{s}}_j\|_2}{\widehat{\rho}_{g,c}}\right), \qquad z_j \overset{\text{iid}}{\sim} \mathcal{N}(0,1), \tag{S33}$$

where $\mathcal{N}(i)$ is a local nearest-neighbor set. For high-resolution data, basis mode instead uses

$$\widetilde{\boldsymbol{\eta}}_g^{\text{para}} = \boldsymbol{\Phi}_g \mathbf{z}, \qquad \mathbf{z} \sim \mathcal{N}(\mathbf{0}, \mathbf{I}_J), \qquad J \ll N, \tag{S34}$$

with basis functions determined by $\widehat{\rho}_{g,c}$. The two constructions reduce per-gene computation from dense GP factorization to local or low-rank operations.

##### S5.3 Blending and variance matching

The pilot-conditioned and parametric fields are combined as

$$\widetilde{\eta}_{ig}^{*} = \lambda_{\text{cond}} \widetilde{\eta}_{ig}^{\text{cond}} + (1 - \lambda_{\text{cond}}) \widetilde{\eta}_{ig}^{\text{para}}, \qquad 0 \le \lambda_{\text{cond}} \le 1. \tag{S35}$$

The blended field is centered and rescaled,

$$\widetilde{\eta}_{ig} = \widehat{\sigma}_{g,c} \frac{\widetilde{\eta}_{ig}^{*} - \overline{\widetilde{\eta}}_{\cdot g}^{*}}{\text{sd}(\widetilde{\boldsymbol{\eta}}_g^{*})}, \tag{S36}$$

so that its marginal variance is $\widehat{\sigma}_{g,c}^2$ irrespective of $\lambda_{\text{cond}}$. Independent nugget noise with variance $\widehat{\tau}_{g,c}^2$ is added afterward. Hyperparameter selection is described in Supplementary Note S7.

##### S5.4 Gene-wise conditional independence and the SaLFC estimand

spaCraft models each gene marginally and does not explicitly reproduce residual gene–gene correlation within domains. Conditional on the generated domain labels, the gene-specific spatial fields and nugget errors are drawn independently, cross-gene dependence therefore arises primarily through the shared tissue architecture and domain-specific mean patterns.

This simplification is aligned with the primary SaLFC endpoint, which is defined by marginal gene-wise rejection probabilities. If $R_g$ denotes rejection of injected gene $g$, then

$$\mathrm{E}\left[\frac{1}{|\mathcal{G}_{\text{spike}}|} \sum_{g \in \mathcal{G}_{\text{spike}}} R_g\right] = \frac{1}{|\mathcal{G}_{\text{spike}}|} \sum_{g \in \mathcal{G}_{\text{spike}}} \Pr(R_g = 1). \tag{S37}$$

Thus, provided the marginal distribution of each gene is calibrated, unmodeled residual gene–gene correlation does not change the expected SaLFC power. It primarily affects dependence among rejections and hence the Monte Carlo variability of their aggregate.

**Algorithm S1** Synthetic sample generation for sample $(k, c)$

**Require:** fitted models $\widehat{M}_1, \widehat{M}_2, \widehat{M}_3$, effect scenario $(p_{\text{target}}, \theta_{\text{spike}})$

Draw the target–reference pair budget and background-domain counts from $\widehat{M}_3$
Set the marginal target share to $\widehat{p}_0$ for controls and $p_{\text{target}}$ for cases, and draw the sample-specific composition
Draw domain centroids and FGKMM parameters from $\widehat{M}_2$; generate $\widetilde{\mathbf{S}}^{(k,c)}$ and generator labels $\widetilde{\mathbf{d}}^{(k,c)}$
**for** each generated gene $g$ **do**
    Draw the sample-specific domain means and sample-level variation from $\widehat{M}_1$
    If $c = 1$ and $g \in \mathcal{G}_{\text{spike}}$, add $\theta_{\text{spike}}$ to the target-domain mean
    Generate the pilot-conditioned and parametric spatial fields and combine them using Eqs. (S32)–(S36)
    Add $\widetilde{\epsilon}_{ig} \sim \mathcal{N}(0, \widehat{\tau}^2_{g,c})$ and form $\widetilde{y}_{ig}$
    Set $\widetilde{X}_{ig} = \max\{\lfloor \exp(\widetilde{y}_{ig}) - 1 \rceil, 0\}$
**end for**
**return** $\widetilde{\mathcal{D}}^{(k,c)} = (\widetilde{\mathbf{X}}^{(k,c)}, \widetilde{\mathbf{S}}^{(k,c)}, \widetilde{\mathbf{d}}^{(k,c)})$

The default Bonferroni procedure also controls family-wise error under arbitrary gene dependence. Explicit multigene dependence modeling would therefore be most relevant for endpoints that depend directly on joint co-expression structure rather than the marginal testing objective considered here.

The generated coordinates remain continuous. Domain recovery is subsequently applied to $(\widetilde{\mathbf{X}}^{(k,c)}, \widetilde{\mathbf{S}}^{(k,c)})$ without access to the generator labels $\widetilde{\mathbf{d}}^{(k,c)}$. Platform-specific coordinate mapping is applied after recovery as described in Supplementary Note S8.

### S6 Pilot-guided domain recovery (pBANKSY)

Within each Monte Carlo replicate, pBANKSY maps $(\widetilde{\mathbf{X}}^{(k,c)}, \widetilde{\mathbf{S}}^{(k,c)})$ to recovered labels $\widehat{\mathbf{d}}^{(k,c)}$ without access to the generator labels $\widetilde{\mathbf{d}}^{(k,c)}$. Building on BANKSY[6], it combines spot-level expression with local spatial context and anchors the resulting embedding to annotated pilot domains. Sample indices are suppressed below for clarity.

#### S6.1 Spatial feature construction

Using genes in $\mathcal{G}_{\text{SVG}}$, let $\mathbf{C}_i$ denote the gene-wise standardized log-expression vector at spot $i$. For its $k$ nearest neighbors $\mathcal{N}_i$, define normalized Gaussian weights

$$w_{ij} \propto \exp\left(-\frac{\|\mathbf{s}_i - \mathbf{s}_j\|_2^2}{\sigma_i^2}\right), \qquad \sum_{j \in \mathcal{N}_i} w_{ij} = 1, \tag{S38}$$

where $\sigma_i$ adapts to the local neighbor distances. The neighborhood mean and local-dispersion blocks are

$$\mathbf{M}_i = \sum_{j \in \mathcal{N}_i} w_{ij}\mathbf{C}_j, \qquad \mathbf{V}_i = \left\{\sum_{j \in \mathcal{N}_i} w_{ij}(\mathbf{C}_j - \mathbf{C}_i)^2\right\}^{1/2}, \tag{S39}$$

with component-wise operations. The mean block represents local expression context, whereas $\mathbf{V}_i$ captures the magnitude of local variation and is typically elevated near domain boundaries.

After gene-wise standardization of the three blocks, let $e_C$, $e_M$, and $e_V$ denote their realized mean squared energies and define

$$\pi_M = \frac{e_M}{e_M + e_V}, \qquad \pi_V = \frac{e_V}{e_M + e_V}.$$

For recovery weight $\lambda_p \in [0, 1]$, the combined feature vector is

$$\mathbf{H}_i = \left(a_C \mathbf{C}_i^\top, a_M \mathbf{M}_i^\top, a_V \mathbf{V}_i^\top\right)^\top, \tag{S40}$$

where

$$a_C = \left(\frac{1-\lambda_p}{e_C}\right)^{1/2}, \qquad a_M = \left(\frac{\lambda_p \pi_M}{e_M}\right)^{1/2}, \qquad a_V = \left(\frac{\lambda_p \pi_V}{e_V}\right)^{1/2}. \tag{S41}$$

Thus, $\lambda_p$ controls the relative contribution of spatial context independently of the realized scales of the feature blocks. Normalization is performed separately for the pilot and synthetic samples.

##### S6.2 Joint embedding and pilot-anchored recovery

Let $\mathbf{H}_p$ and $\widetilde{\mathbf{H}}$ denote the pilot and synthetic feature matrices, each with $3|\mathcal{G}_{\text{SVG}}|$ columns. A single PCA is fitted to their joint matrix,

$$\begin{pmatrix} \mathbf{U}_p \\ \mathbf{U}_{\text{syn}} \end{pmatrix} = \text{PCA}\left(\begin{pmatrix} \mathbf{H}_p \\ \widetilde{\mathbf{H}} \end{pmatrix}\right). \tag{S42}$$

The annotated pilot centroid for domain $d$ in the retained PCA space is

$$\mathbf{q}_d = \frac{1}{n_{p,d}} \sum_{i:d_{p,i}=d} \mathbf{U}_{p,i}. \tag{S43}$$

By default, $k$-means with $D = |\mathcal{A}|$ clusters is applied to $\mathbf{U}_{\text{syn}}$ using $\{\mathbf{q}_d : d \in \mathcal{A}\}$ as initial centers. Because cluster indices are arbitrary, recovered clusters are matched to pilot domains by

$$\widehat{\pi} = \operatorname*{argmin}_{\pi \in \mathfrak{S}_D} \sum_{v=1}^{D} \left\|\widehat{\mathbf{q}}_v^{\text{syn}} - \mathbf{q}_{\pi(v)}\right\|_2^2, \qquad \widehat{d}_i = \widehat{\pi}(d_i^*), \tag{S44}$$

using the Hungarian algorithm. This assigns biological domain identities without altering the recovered partition and without using $\widetilde{\mathbf{d}}$. An optional stricter mode assigns each synthetic spot directly to its nearest pilot-domain centroid in the joint embedding. For fixed neighborhood size, PCA dimension, and number of domains, recovery scales approximately linearly with the number of synthetic spots. Pilot feature blocks are computed once and reused across Monte Carlo replicates.

#### S7 Weighting scheme

The recovery weight $\lambda_p$ and texture weight $\lambda_{\text{cond}}$ control distinct parts of the generate–recover–test loop and are tuned sequentially before the final power calculation. The first

**Algorithm S2** Pilot-guided domain recovery (pBANKSY)

**Require:** synthetic sample $(\widetilde{\mathbf{X}}, \widetilde{\mathbf{S}})$, annotated pilot reference, recovery weight $\lambda_p$, neighbor count $k$
1: Construct standardized expression, neighborhood-mean, and local-dispersion blocks for pilot and synthetic samples
2: Normalize block energies and assemble $\mathbf{H}_p$ and $\widetilde{\mathbf{H}}$
3: Obtain a joint PCA embedding and compute pilot-domain centroids
4: Run $k$-means on the synthetic embedding initialized at the pilot centroids
5: Match recovered clusters to pilot domains using the Hungarian assignment
6: **return** recovered labels $\widehat{\mathbf{d}}^{(k,c)}$

stage tunes domain recovery from annotated pilot samples; conditional on this choice, the second stage tunes synthetic spatial texture for each endpoint and effect size.

##### S7.1 Stage 1: recovery weight

The recovery weight $\lambda_p$ controls the contribution of spatial context to the pBANKSY embedding. It is selected by pilot cross-validation. In each split $b$, one pilot sample is treated as the query and same-group samples form the annotated reference; the query labels are withheld during recovery and used only for evaluation. For candidate grid $\Lambda_p$,

$$\widehat{\lambda}_p = \underset{\lambda \in \Lambda_p}{\operatorname{argmax}} \frac{1}{B_{\text{cv}}} \sum_{b=1}^{B_{\text{cv}}} \text{ARI}\left(\widehat{\mathbf{d}}_{\lambda}^{(b)}, \mathbf{d}^{(b)}\right). \tag{S45}$$

The selected $\widehat{\lambda}_p$ is then fixed across endpoints and effect sizes.

##### S7.2 Stage 2: texture weight

Conditional on $\widehat{\lambda}_p$, the texture weight $\lambda_{\text{cond}}$ balances pilot-conditioned and parametric spatial structure. Let $e \in \{SaLFC, LOR\}$ denote the endpoint and let $\theta_e$ denote its effect parameter: $\theta_{SaLFC} = \theta_{\text{spike}}$ and $\theta_{LOR} = p_{\text{target}}$. For tuning replicate $r$, define

$$T_{SaLFC,r} = \frac{1}{|\mathcal{G}_{\text{spike}}|} \sum_{g \in \mathcal{G}_{\text{spike}}} |T_{g,r}|, \qquad T_{LOR,r} = T_{\text{comp},r}. \tag{S46}$$

For candidate $\lambda \in \Lambda_c$, the endpoint signal is summarized by the regularized noncentrality

$$\widehat{\delta}_e(\lambda \mid \theta_e) = \frac{\overline{T}_e(\lambda \mid \theta_e)}{\text{sd}_r\{T_{e,r}(\lambda \mid \theta_e)\} + \varepsilon s_{0,e}}, \tag{S47}$$

where $s_{0,e}$ is a pooled scale and $\varepsilon > 0$ stabilizes the denominator. This criterion favors spatial textures that produce a strong and reproducible endpoint signal rather than low variance alone.

Each candidate is also evaluated under the corresponding null: $\theta_{\text{spike}} = 0$ for SaLFC and $p_{\text{target}} = \widehat{p}_0$ for LOR. Let $\widehat{\alpha}_e(\lambda)$ denote its empirical null rejection rate. To discourage

**Algorithm S3** Two-stage tuning of $(\lambda_p, \lambda_{\text{cond}})$

**Require:** annotated pilot, fitted models, grids $\Lambda_p$ and $\Lambda_c$, endpoint-specific effect grids

```
Select \hat{λ}_p by pilot cross-validation using Eq. (S45)
for each endpoint e and effect θ_e do
    for λ ∈ Λ_c do
        Generate tuning cohorts and compute \hat{δ}_e(λ | θ_e)
        Generate matched-null cohorts and estimate \hat{α}_e(λ)
        Compute the calibration weight w_e(λ)
    end for
    Select \hat{λ}_{cond,e}(θ_e) by Eq. (S49)
end for
return \hat{λ}_p and {\hat{λ}_{cond,e}(θ_e)}
```

anti-conservative choices, define

$$w_e(\lambda) = \exp\left[-\frac{1}{2}\left\{\frac{[\widehat{\alpha}_e(\lambda) - \alpha]_+}{\alpha(\kappa - 1)}\right\}^2\right], \qquad \kappa > 1. \tag{S48}$$

The endpoint- and effect-specific texture weight is then

$$\widehat{\lambda}_{\text{cond},e}(\theta_e) = \operatorname*{argmax}_{\lambda \in \Lambda_c} \widehat{\delta}_e(\lambda \mid \theta_e) w_e(\lambda). \tag{S49}$$

Thus, tuning favors a large, stable endpoint signal while penalizing candidate generators that inflate null rejection. The selected weights are fixed before the corresponding final power simulations.

### S8 Coordinate rearrangement and optimal transport

After domain recovery, the continuous synthetic coordinates are rearranged onto the lattice of a randomly selected same-group pilot sample. Expression and recovered labels remain fixed, only the coordinates used by the downstream spatial analysis are updated. We write $\widetilde{\mathbf{S}}^{\text{cont}}$ for the continuous coordinates before rearrangement and $\widetilde{\mathbf{S}}$ for the final lattice-mapped coordinates. For recovered domain $d$, let

$$\widetilde{\mathbf{S}}_d^{\text{cont}} = \left\{\widetilde{\mathbf{s}}_i^{\text{cont}} : \widehat{d}_i = d\right\}, \qquad \widehat{n}_d = \left|\widetilde{\mathbf{S}}_d^{\text{cont}}\right|,$$

and let $\mathbf{P}_d$ denote the lattice sites annotated as domain $d$ in the selected pilot sample, with $n_{p,d} = |\mathbf{P}_d|$.

#### S8.1 Composition-aware reference territories

When the target share differs from the pilot composition, the target and reference territories are adjusted before transport. Let $\mathcal{P} = \mathbf{P}_T \cup \mathbf{P}_R$ and define

$$s(\mathbf{p}) = d(\mathbf{p}, \mathbf{P}_T) - d(\mathbf{p}, \mathbf{P}_R), \qquad d(\mathbf{p}, \mathbf{P}_d) = \min_{\mathbf{q} \in \mathbf{P}_d} \|\mathbf{p} - \mathbf{q}\|_2. \tag{S50}$$

For group $c$, let

$$p_c^{\text{gen}} = \begin{cases} \widehat{p}_0, & c = 0, \\ p_{\text{target}}, & c = 1. \end{cases}$$

The $\lfloor p_c^{\text{gen}} |\mathcal{P}| \rceil$ sites with the smallest scores are assigned to the target territory and the remainder to the reference territory. Thus, expansion and contraction proceed primarily along the target–reference boundary.

For each recovered domain, a reference set $\mathbf{R}_d$ of size $\widehat{n}_d$ is drawn from the corresponding pilot territory. Sampling is without replacement when possible; if $\widehat{n}_d > n_{p,d}$, additional sites are sampled with replacement and given a small spatial jitter to avoid exact coordinate duplication.

##### S8.2 Gaussian transport and lattice assignment

Let $(\boldsymbol{\mu}_S, \boldsymbol{\Sigma}_S)$ and $(\boldsymbol{\mu}_R, \boldsymbol{\Sigma}_R)$ denote the empirical moments of $\widetilde{\mathbf{S}}_d^{\text{cont}}$ and $\mathbf{R}_d$, respectively. The Gaussian transport map is

$$\mathcal{T}_d(\mathbf{s}) = \boldsymbol{\mu}_R + \mathbf{A}_d(\mathbf{s} - \boldsymbol{\mu}_S), \qquad \mathbf{A}_d = \boldsymbol{\Sigma}_S^{-1/2} \left( \boldsymbol{\Sigma}_S^{1/2} \boldsymbol{\Sigma}_R \boldsymbol{\Sigma}_S^{1/2} \right)^{1/2} \boldsymbol{\Sigma}_S^{-1/2}. \tag{S51}$$

This aligns the location, scale, and orientation of the generated domain with the pilot territory while preserving its relative internal geometry. The transported spots are then assigned bijectively to $\mathbf{R}_d$ using a nearest-unoccupied search over local lattice candidates. Repeating this procedure across recovered domains yields the final coordinates $\widetilde{\mathbf{S}}^{(k,c)}$. Because $\widetilde{\mathbf{X}}^{(k,c)}$ and $\widehat{\mathbf{d}}^{(k,c)}$ are unchanged, rearrangement preserves the recovered domain counts and expression contrasts while adapting the spatial layout to the measurement platform.

**Algorithm S4** Coordinate rearrangement and lattice assignment

**Require:** continuous coordinates $\widetilde{\mathbf{S}}^{\text{cont}}$, recovered labels $\widehat{\mathbf{d}}$, same-group pilot pool, scenario target share $p_c^{\text{gen}}$

1: Draw one same-group pilot sample as the lattice reference
2: If needed, update the target and reference territories using Eq. (S50)
3: **for** each recovered domain $d$ **do**
4: Form $\widetilde{\mathbf{S}}_d^{\text{cont}}$ and a count-matched reference set $\mathbf{R}_d$
5: Apply the Gaussian transport map in Eq. (S51)
6: Assign transported spots to available sites in $\mathbf{R}_d$
7: **end for**
8: **return** final lattice coordinates $\widetilde{\mathbf{S}}^{(k,c)}$

#### S9 Validity of the endpoint tests

Both endpoints reduce each biological sample to a recovered-domain contrast. Let $Z^{(k,c)}$ denote the sample-level contrast and $\theta_0$ a pilot-derived center. The generic estimand is

$$\theta = \mathrm{E}\left(Z^{(k,1)}\right) - \mathrm{E}\left(Z^{(k,0)}\right) - \theta_0, \tag{S52}$$

with estimator $\widehat{\theta} = \bar{Z}_1 - \bar{Z}_0 - \theta_0$. Motivated by TREAT[7], for margin $\gamma \geq 0$, we test

$$H_0 : \theta \leq \gamma \qquad \text{versus} \qquad H_1 : \theta > \gamma$$

using

$$T = \frac{\widehat{\theta} - \gamma}{\widehat{s}}, \tag{S53}$$

where $\widehat{s}^2$ consistently estimates $\mathrm{Var}(\widehat{\theta})$. For SaLFC, $\theta_0 = 0$ and $\gamma = \gamma_g$; for LOR, $\theta_0 = \widehat{\beta}_1$ and $\gamma = \gamma_{\text{comp}}$. All statements below are conditional on the pilot, so fitted models, tuning parameters, centers, and margins are treated as fixed.

**Assumption S1** (Regularity). *As $K_0, K_1 \to \infty$, $K_1/(K_0 + K_1) \to \omega \in (0, 1)$. Within each group, the sample-level contrasts are independent and identically distributed with finite $(2 + \xi)$th moment and nonzero finite variance. Further, $\widehat{s}^2/\,Var(\widehat{\theta}) \xrightarrow{p} 1$.*

**Theorem S1** (Asymptotic validity). *Under Assumption S1, $T \xrightarrow{d} \mathcal{N}(0, 1)$ at $\theta = \gamma$. Hence the test rejecting for $T > z_{1-\alpha}$ has asymptotic level $\alpha$ over $H_0 : \theta \le \gamma$ and is consistent for every fixed $\theta > \gamma$.*

*Proof Sketch.* The two-sample central limit theorem gives asymptotic normality of $\widehat{\theta}$. At the boundary $\theta = \gamma$, consistency of $\widehat{s}^2$ and Slutsky's theorem yield the standard normal limit. For $\theta < \gamma$ the rejection probability is asymptotically no larger than at the boundary, whereas for $\theta > \gamma$, $(\widehat{\theta} - \gamma)/\widehat{s} \to \infty$ in probability. □

The theorem applies to contrasts defined by the recovered labels $\widehat{\mathbf{d}}$ obtained by pBANKSY. Recovery error may attenuate the available effect by mixing target and reference spots, but does not invalidate the test when the resulting sample-level contrasts satisfy the stated regularity conditions. Thus, inference targets the effect available to the downstream analysis rather than the latent effect defined by $\widetilde{\mathbf{d}}$. Additionally, the provided software package includes the Wilcoxon rank-sum test as an optional non-parametric alternative.

#### S9.1 Endpoint I: SaLFC

For gene $g$, define

$$Z_g^{(k,c)} = \bar{y}_{g,T}^{(k,c)} - \bar{y}_{g,R}^{(k,c)},$$

where the means are computed using recovered labels. Let

$$\widehat{\mathcal{I}}_d^{(k,c)} = \{i : \widehat{d}_i = d\}, \qquad \widehat{n}_d^{(k,c)} = |\widehat{\mathcal{I}}_d^{(k,c)}|.$$

At the final rearranged coordinates, the fitted spot-level covariance is

$$\widehat{\Omega}_g^{(k,c)}(i,j) = \widehat{\sigma}_{g,c}^2 \exp\left( -\frac{\|\widetilde{\mathbf{s}}_i^{(k,c)} - \widetilde{\mathbf{s}}_j^{(k,c)}\|_2}{\widehat{\rho}_{g,c}} \right) + \widehat{\tau}_{g,c}^2 \mathbb{1}\{i = j\}. \tag{S54}$$

The resulting within-sample variance of the target–reference contrast is

$$v_g^{(k,c)} = \frac{1}{(\widehat{n}_T^{(k,c)})^2} \sum_{i,j \in \widehat{\mathcal{I}}_T^{(k,c)}} \widehat{\Omega}_g^{(k,c)}(i,j) + \frac{1}{(\widehat{n}_R^{(k,c)})^2} \sum_{i,j \in \widehat{\mathcal{I}}_R^{(k,c)}} \widehat{\Omega}_g^{(k,c)}(i,j) - \frac{2}{\widehat{n}_T^{(k,c)} \widehat{n}_R^{(k,c)}} \sum_{i \in \widehat{\mathcal{I}}_T^{(k,c)}} \sum_{j \in \widehat{\mathcal{I}}_R^{(k,c)}} \widehat{\Omega}_g^{(k,c)}(i,j). \tag{S55}$$

The between-sample biological contrast variance from $\widehat{M}_1$ is

$$\widehat{V}_{\text{bio},g,c} = 2\,(1\widehat{\varrho}_{TR,g,c})\,\widehat{\sigma}^2_{\text{bio},g,c}. \tag{S56}$$

Because pilot sample sizes are small, these variances are moderated across genes,

$$\widetilde{V}_{\text{bio},g,c} = \frac{d_{0,c}V_{0,c} + (K_c - 1)\widehat{V}_{\text{bio},g,c}}{d_{0,c} + K_c - 1}, \tag{S57}$$

following empirical-Bayes variance moderation[8]. With $\bar{v}_{g,c} = K_c^{-1}\sum_k v_g^{(k,c)}$, the variance of the group contrast is estimated by

$$\widehat{s}_g^2 = \sum_{c\in\{0,1\}} \frac{\widetilde{V}_{\text{bio},g,c} + \bar{v}_{g,c}}{K_c}. \tag{S58}$$

A Satterthwaite approximation provides the corresponding degrees of freedom, and the statistic in Methods equation (20) is referred to the resulting $t$ distribution. Bonferroni adjustment controls FWER under arbitrary dependence among genes. BH (Benjamini-Hochberg) adjustment is also reported for FDR control under its standard dependence conditions.

#### S9.2 Endpoint II: LOR

For LOR, sample $(k, c)$ contributes the recovered-domain log odds

$$L^{(k,c)} = \log\left(\frac{\widehat{n}_T^{(k,c)} + h}{\widehat{n}_R^{(k,c)} + h}\right), \qquad h > 0, \tag{S59}$$

where $h$ is a continuity correction with 0.5 as default. The baseline-anchored estimand is

$$\Delta = \mathrm{E}\left(L^{(k,1)}\right) - \mathrm{E}\left(L^{(k,0)}\right) - \widehat{\beta}_1. \tag{S60}$$

Conditional on the pilot, $\widehat{\beta}_1$ is fixed. If $\widehat{\sigma}^2_{L,c}$ is the sample variance of $\{L^{(k,c)}\}_{k=1}^{K_c}$, the Welch variance estimator is

$$\widehat{s}^2_{\text{LOR}} = \frac{\widehat{\sigma}^2_{L,1}}{K_1} + \frac{\widehat{\sigma}^2_{L,0}}{K_0}. \tag{S61}$$

The statistic in Methods equation (24) is referred to a Welch–Satterthwaite $t$ distribution. Spatial organization enters through the recovered domain counts and therefore requires no additional spot-level variance term.

#### S9.3 Bootstrap-calibrated margins

The margins $\gamma_g$ and $\gamma_{\text{comp}}$ are calibrated once from pilot-based null simulations and then held fixed for all candidate sample sizes. Null cohorts are generated with $\theta_{\text{spike}} = 0$ for SaLFC and $p_{\text{target}} = \widehat{p}_0$ for LOR, and pass through the same generate–recover–rearrange–test pipeline as the power simulations.

For null replicate $b = 1, \ldots, B_0$, let $\widehat{\delta}^0_{g,b}$ and $\widehat{\Delta}^0_b$ denote the resulting endpoint estimates.

The margins are

$$\gamma_g = Q_{0.50}\left(\left\{|\widehat{\delta}^0_{g,b}|\right\}_{b=1}^{B_0}\right), \qquad \gamma_{\text{comp}} = Q_{0.50}\left(\left\{|\widehat{\Delta}^0_b|\right\}_{b=1}^{B_0}\right). \tag{S62}$$

Hence rejection requires an effect exceeding the typical null fluctuation, rather than merely a nonzero estimate. Because these margins are fixed before evaluating candidate $K$, differences among power curves reflect sample size rather than a changing rejection boundary.

### S10 Held-out validation

Held-out validation assessed whether sample sizes predicted from a small pilot agree with those obtained from independent real samples. For each of $B = 100$ repeated splits, $K_{\text{pilot}}$ samples per group were used exclusively to construct the gene sets, fit $\widehat{M}_1, \widehat{M}_2, \widehat{M}_3$, tune the recovery and texture weights, and calibrate the testing margins. The remaining samples formed the held-out cohort.

For the 5xFAD data, the full cohort contained 8 control and 12 case samples, with $K_{\text{pilot}} = 3$ per group. Each split therefore left 5 control and 9 case samples. Pilot-based synthetic power was evaluated for $K \in \{3, 4, 5, 6\}$, whereas held-out validation was restricted to $K \in \{3, 4, 5\}$ by the smaller control arm.

#### S10.1 Synthetic and held-out validation arms

The pilot-simulated arm followed the complete generate–recover–rearrange–test pipeline and produced $\widehat{\pi}_e^{\text{sim},(b)}(K)$. For the held-out arm, the identical endpoint-specific effect was applied to balanced subsets of independent held-out samples. Subsequent domain recovery and endpoint testing yielded $\widehat{\pi}_e^{\text{HO},(b)}(K)$, deliberately bypassing the model fitting and data generation steps. Crucially, the held-out annotations served exclusively to impose this effect. Both arms were then evaluated uniformly using the labels recovered by pBANKSY.

For SaLFC, injected genes in the annotated case target domain were shifted by

$$y_{ig}^{(k,1)} \longmapsto y_{ig}^{(k,1)} + \theta_{\text{spike}} \mathbb{1}\{d_i = T,\, g \in \mathcal{G}_{\text{spike}}\}. \tag{S63}$$

For LOR, a target share with marginal mean $p_{\text{target}}$ was drawn using the fitted logit-normal variation $\widehat{\sigma}_{\text{re}}$, while preserving each sample's observed target–reference budget. The resulting change in the target count was achieved by relabelling the annotated spots that were geometrically closest to the target-reference interface. Expression at reassigned spots was redrawn from $\widehat{M}_1$ using the parameters of the new domain, preserving consistency between composition and the expression pattern presented to pBANKSY.

#### S10.2 Power and sample-size agreement

At the common sample sizes $K \in \{3, 4, 5\}$, agreement between the pilot-simulated and held-out power estimates was summarized by

$$\Delta\pi_e^{(b)}(K) = \widehat{\pi}_e^{\text{HO},(b)}(K) - \widehat{\pi}_e^{\text{sim},(b)}(K). \tag{S64}$$

Median differences remained within 0.034 of zero for every endpoint and sample size. Held-out estimates were more variable, reflecting the finite subsets of independent real samples available in each split.

| Endpoint | $K$ | Median $\Delta\pi$ | Mean $\Delta\pi$ | IQR of $\Delta\pi$ | Median $\lvert\Delta\pi\rvert$ |
|---|---|---|---|---|---|
| SaLFC | 3 | −0.034 | −0.073 | [−0.256, +0.140] | 0.175 |
| | 4 | −0.007 | −0.108 | [−0.240, +0.068] | 0.110 |
| | 5 | −0.020 | −0.125 | [−0.223, +0.029] | 0.103 |
| LOR | 3 | −0.017 | −0.049 | [−0.233, +0.108] | 0.167 |
| | 4 | 0.000 | +0.017 | [−0.100, +0.142] | 0.133 |
| | 5 | 0.000 | +0.022 | [−0.100, +0.108] | 0.100 |

**Supplementary Table S1 | Held-out validation of power at fixed sample sizes.** Paired held-out minus pilot-simulated power differences over $B = 100$ splits, with $N_{\text{rep}} = 30$ replicate cohorts per arm. Comparisons are restricted to $K \in \{3, 4, 5\}$, which are supported by both validation arms.

We next compared the sample-size recommendations derived from the two power curves. For each split and endpoint, the simulated and held-out power values were fitted by monotone shape-constrained curves. Let $\kappa_e^{s,(b)}$ denote the crossing of target power $\pi^*$ for $s \in \{\text{sim}, \text{HO}\}$. We define

$$K^{*,(b)}_{\text{sim},e} = \left\lceil \kappa^{(b)}_{\text{sim},e} \right\rceil, \qquad K^{*,(b)}_{\text{HO},e} = \left\lfloor \kappa^{(b)}_{\text{HO},e} \right\rfloor. \tag{S65}$$

The prospective recommendation is rounded upward, whereas the held-out crossing is treated as an empirical estimate. Comparisons were retained when both recommendations were supported by the held-out range $K \leq 5$. Their signed difference was

$$\Delta K^{(b)}_e = K^{*,(b)}_{\text{HO},e} - K^{*,(b)}_{\text{sim},e}, \tag{S66}$$

so $\Delta K^{(b)}_e > 0$ indicates underestimation by the pilot-based design.

Both recommendations were evaluable in 65 of 100 splits for SaLFC and 71 of 100 splits for LOR, with the remaining splits censored because one or both power curves did not cross the target within the supported sample-size range. The mean signed errors were −0.26 samples (90% CI, $[-0.43, -0.09]$) for SaLFC and 0.13 samples (90% CI, $[-0.07, 0.32]$) for LOR, with median error zero for both. When restricting the analysis to evaluable splits (i.e., omitting censored outcomes), the absolute error did not exceed a single sample in 60/65 (92%) and 63/71 (89%) of the partitions, respectively. Two one-sided equivalence tests with a prespecified margin of one sample confirmed that the mean signed errors lay within the equivalence bounds of −1 and +1 sample ($p < 10^{-9}$ for both endpoints).

| Endpoint | $\leq 3$ | 4 | 5 | $> 5$ |
|---|---|---|---|---|
| SaLFC | 44 | 36 | 14 | 6 |
| LOR | 46 | 22 | 26 | 6 |

**Supplementary Table S2 | Distribution of pilot-simulated sample-size recommendations.** Number of the $B = 100$ pilot splits yielding each $K^*_{\text{sim},e}$. The effect scenario was fixed across splits within each endpoint. The first category includes crossings at or below the smallest evaluated size, and the final category includes recommendations above the held-out support.

**Algorithm S5** Held-out validation by repeated splitting

**Require:** full cohort, pilot size $K_{\text{pilot}}$, number of splits $B$, effect scenario, target power $\pi^*$

**for** $b = 1, \ldots, B$ **do**
  Split each group into pilot and held-out samples
  Using only the pilot, construct gene sets, fit $\widehat{M}_1, \widehat{M}_2, \widehat{M}_3$, tune weights, and calibrate testing margins
  Estimate pilot-simulated power by the complete generate–recover–rearrange–test pipeline
  Apply the same effect to balanced held-out subsets, recover domains, and estimate held-out power
  Fit monotone power curves and obtain $K^*_{\text{sim},e}$ and $K^*_{\text{HO},e}$
  Retain the split when both recommendations lie within the common held-out support
**end for**
**return** $\Delta K_e = K^*_{\text{HO},e} - K^*_{\text{sim},e}$ and fixed-$K$ power differences

### Supplementary Figures

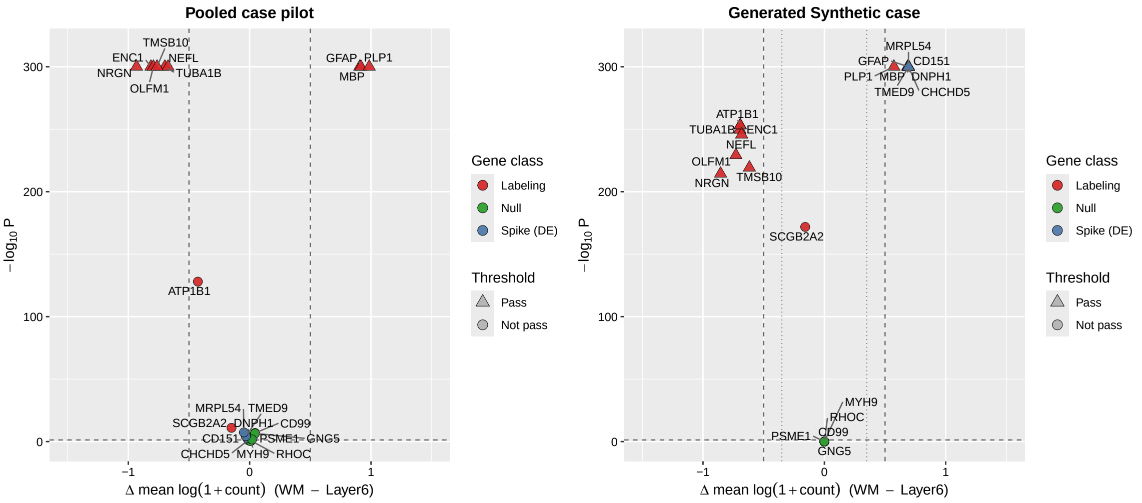

**Supplementary Fig. S1 | Injected SaLFC effects remain distinguishable after domain recovery** (DLPFC, white matter versus Layer 6). Genes are shown by target-minus-reference contrast and $-\log_{10} P$, with colors indicating $\mathcal{G}_{\text{spike}} \subset \mathcal{G}_{\text{Null}} \subset \mathcal{G}_{\text{SVG}}$. In the pilot (left), injection genes have small baseline contrasts before perturbation. In a synthetic case evaluated using recovered labels $\widehat{\mathbf{d}}$ (right), the injected genes *(MRPL54, DNPH1, CD151, TMED9, CHCHD5)* separate from the noninjected null genes, which remain near zero.

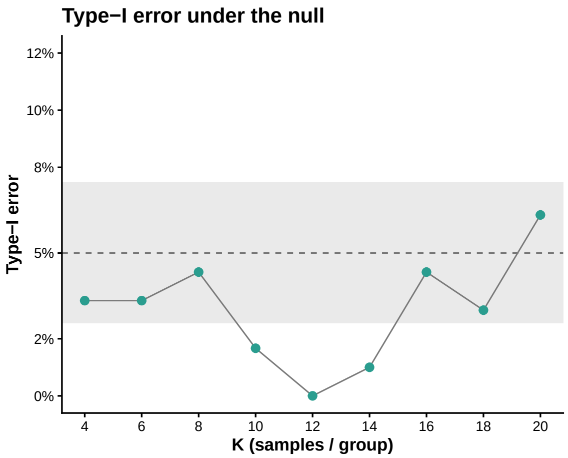

**Supplementary Fig. S2 | Null calibration of the composition endpoint.** Results are shown for the DLPFC Visium pilot with white matter versus Layer 6 as the target and reference pair. Empirical LOR rejection rates against the sample size per group $K$ under the compositional null of the deployed pipeline (fitted case models, target share $\widehat{p}_0$, labels recovered by pBANKSY). Domain recovery induces an offset near 0.08 in the recovered log odds. The fixed margin $\gamma_{\mathrm{comp}} = 0.15$, the median absolute null statistic of the full pipeline at $K_{\mathrm{ref}} = 8$ on disjoint replicates, exceeds this offset, so the generated null lies within $H_0 : \Delta \leq \gamma_{\mathrm{comp}}$ and Theorem S1 applies. Points, 300 Monte Carlo replicates each. Dashed line, $\alpha = 0.05$. Gray band, its 95% binomial interval. All rates lie within or below the band.

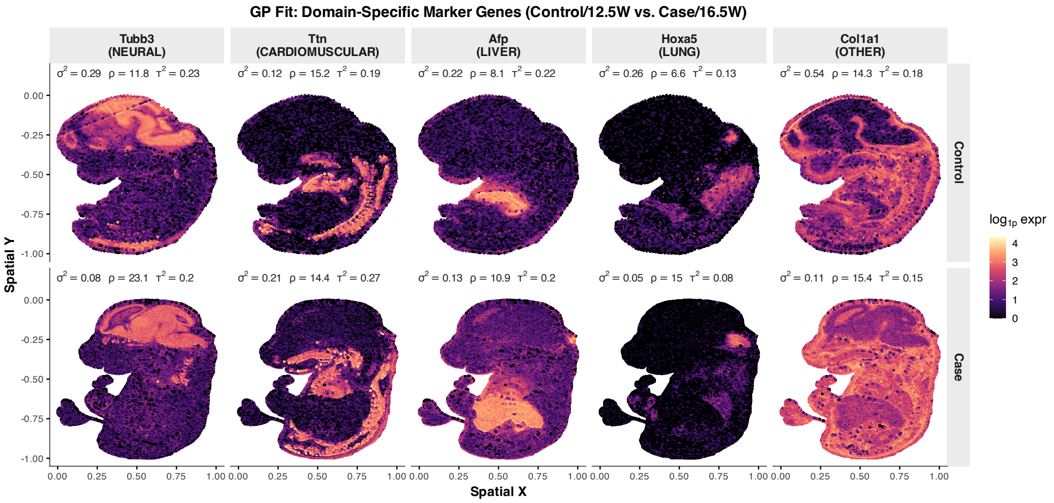

**Supplementary Fig. S3 | Expression-model fits for domain markers in the mouse-embryo Stereo-seq cohort.** Spatial log-expression of one marker per consolidated anatomical domain in one E12.5 control (top) and one E16.5 case (bottom) sample, with fitted Gaussian-process partial sill $\sigma^2$, range $\rho$, and nugget $\tau^2$ shown above each panel. Expression is shown on pseudo-spots formed by domain-stratified nearest-neighbor binning (Methods), so the fitted range is defined on the pseudo-spot lattice. Color scale, $\log(1 + \text{count})$.

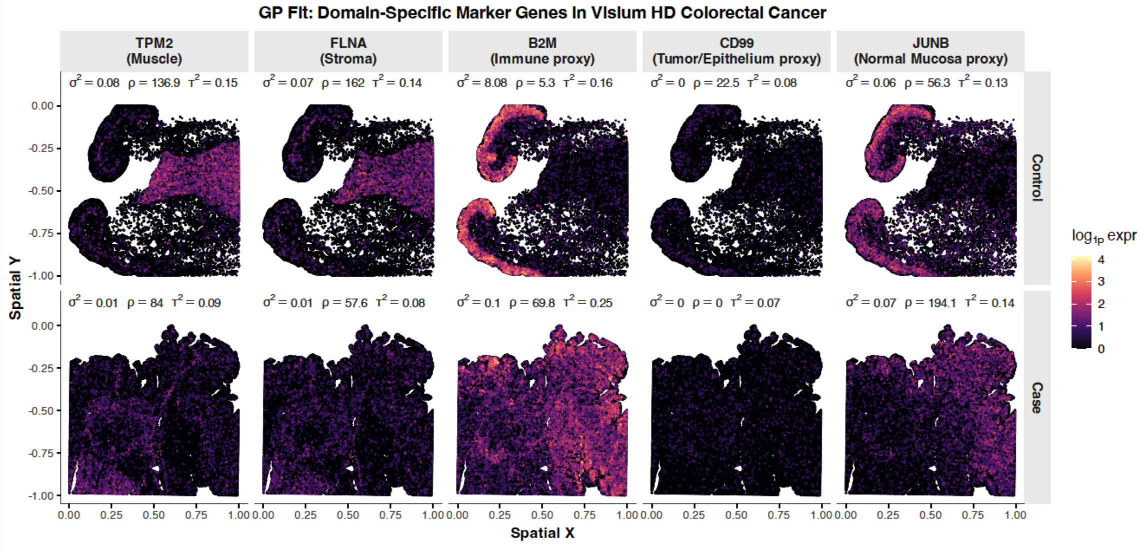

**Supplementary Fig. S4 | Expression-model fits for domain markers in the colorectal cancer Visium HD cohort.** Spatial log-expression of one marker per histological domain in one normal-adjacent control (top) and one tumor case (bottom) sample, with fitted Gaussian-process partial sill $\sigma^2$, range $\rho$, and nugget $\tau^2$ shown above each panel. Spatial signal was weaker for several markers in the tumor sample, including *TPM2* ($\sigma^2$: 0.08 to 0.01) and *FLNA* (0.07 to 0.01), and this difference was retained by the fitted model. Color scale, $\log(1 + \text{count})$.

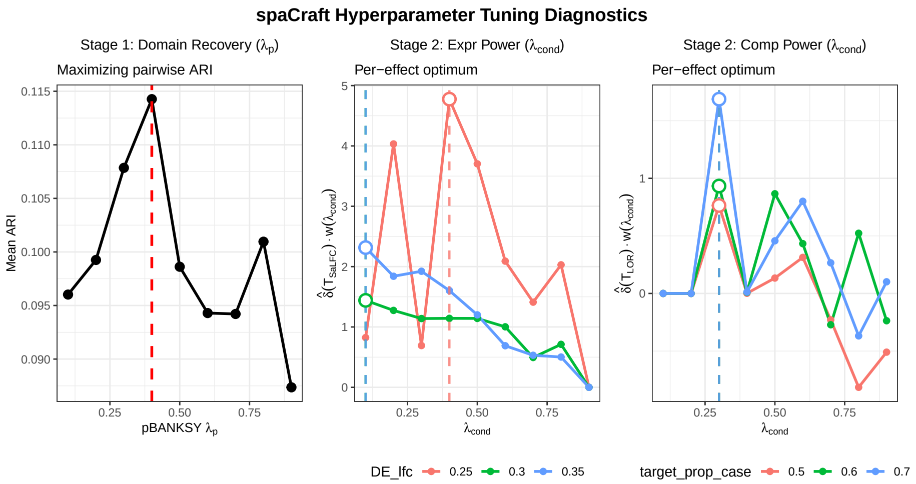

**Supplementary Fig. S5 | Two-stage tuning diagnostics.** Stage 1 selects the recovery weight $\lambda_p$ by mean adjusted Rand index between recovered partitions and withheld pilot annotations (Eq. (S45)). Conditional on this value, Stage 2 selects the endpoint- and effect-specific texture weight $\lambda_{\text{cond}}$ by regularized noncentrality (Eq. (S47)) with a null-calibration guard. Results are shown for the DLPFC pilot (Supplementary Note S7).

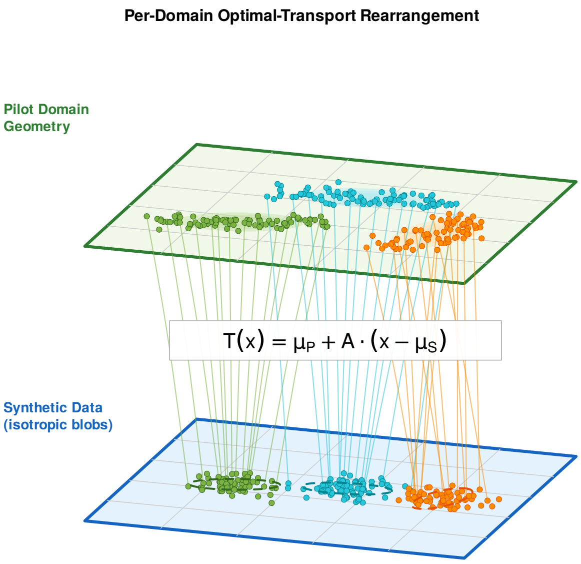

**Supplementary Fig. S6 | Per-domain Gaussian optimal-transport alignment.** Visual summary of Algorithm S4. Each recovered synthetic domain is mapped to a count-matched set of lattice sites derived from the corresponding domain of a same-group pilot sample, using a Gaussian affine transport followed by bijective nearest-site assignment.

#### Supplementary Tables

| # | Gene | $\bar{y}_T$ | $\bar{y}_R$ | $\Delta_g^{TR}$ |
|---|---|---|---|---|
| **$\mathcal{G}_{\mathrm{SVG}}$, labeling set (domain recovery),** $\lvert\mathcal{G}_{\mathrm{SVG}}\rvert = 21$ | | | | |
| Labeling-only genes ($\mathcal{G}_{\mathrm{SVG}} \setminus \mathcal{G}_{\mathrm{Null}}$, $n = 11$) | | | | |
| *GFAP*, *TUBA1B*, *NRGN*, *ENC1*, *OLFM1*, *ATP1B1*, *SCGB2A2*, *NEFL*, *TMSB10*, *MBP*, *PLP1* | | | | |
| **$\mathcal{G}_{\mathrm{Null}}$, empirical-null set (type-I calibration),** $\lvert\mathcal{G}_{\mathrm{Null}}\rvert = 10$ | | | | |
| $\mathcal{G}_{\mathrm{spike}}$, injection set (SaLFC effect), $n = 5$ | | | | |
| 1 | *MRPL54* | 0.1157 | 0.1868 | −0.0710 |
| 2 | *DNPH1* | 0.0775 | 0.1430 | −0.0655 |
| 3 | *CD151* | 0.1072 | 0.1567 | −0.0495 |
| 4 | *TMED9* | 0.0837 | 0.1228 | −0.0391 |
| 5 | *CHCHD5* | 0.0860 | 0.1183 | −0.0323 |
| Noninjected null genes ($\mathcal{G}_{\mathrm{Null}} \setminus \mathcal{G}_{\mathrm{spike}}$, $n = 5$) | | | | |
| 6 | *RHOC* | 0.1263 | 0.1448 | −0.0185 |
| 7 | *GNG5* | 0.1423 | 0.1573 | −0.0150 |
| 8 | *MYH9* | 0.1547 | 0.1417 | +0.0130 |
| 9 | *PSME1* | 0.1337 | 0.1254 | +0.0083 |
| 10 | *CD99* | 0.1454 | 0.1385 | +0.0069 |

**Supplementary Table S3 | Gene sets and target–reference statistics for the DLPFC Visium analysis** (reference pilot sample 151673; $(T, R)$ = white matter versus Layer 6). The sets are nested, $\mathcal{G}_{\mathrm{spike}} \subset \mathcal{G}_{\mathrm{Null}} \subset \mathcal{G}_{\mathrm{SVG}}$. $\mathcal{G}_{\mathrm{SVG}}$ is used for domain recovery, $\mathcal{G}_{\mathrm{Null}}$ for SaLFC testing, and $\mathcal{G}_{\mathrm{spike}}$ receives the imposed expression effect. Injection genes are the null genes with the largest baseline $|\Delta_g^{TR}|$. For each null gene, $\bar{y}_T$ and $\bar{y}_R$ are the mean $\log(1 + \text{count})$ expression in WM and Layer 6, with $\Delta_g^{TR} = \bar{y}_T - \bar{y}_R$. Labeling-only genes are listed by name.

| Set | Stereo-seq mouse embryo (lung vs neural) | Visium HD colorectal cancer (normal epithelium vs stroma) |
|---|---|---|
| $\mathcal{G}_{\mathrm{SVG}}$ labeling set (domain recovery) | $n = 40$. Labeling-only genes ($\mathcal{G}_{\mathrm{SVG}} \setminus \mathcal{G}_{\mathrm{Null}}$, $n = 30$) *Ttn, Tnni1, Tnnc1, Myl1, Actc1, Tnnt1, Afp, Alb, Mt2, Car2, Apoa2, Slc25a37, Hoxa5, Unc5c, Pde5a, Ror1, Adam12, Tbx2, Tubb3, Rtn1, Crmp1, Fez1, Fabp7, Map2, Gpc3, Col1a2, Col1a1, Col3a1, Cnn2, Pdgfra* | $n = 47$. Labeling-only genes ($\mathcal{G}_{\mathrm{SVG}} \setminus \mathcal{G}_{\mathrm{Null}}$, $n = 37$) *PIGR, MT-CO3, MT-ATP6, MT-CO2, MT-ND4, IGKC, PHGR1, MT-CYB, TSPAN8, MT-ND3, JCHAIN, VIM, TMSB4X, EGR1, TXNDC5, B2M, FCGBP, MT-ND4L, SELENOP, RNASE1, CD74, FTL, MCL1, ZFP36, DES, TAGLN, MYL9, MYH11, ACTG2, CSRP1, TPM2, TPM1, FLNA, CNN1, A2M, JUNB, ADAMTS1* |
| $\mathcal{G}_{\mathrm{Null}}$ empirical-null set (type-I calibration) | $n = 10$. Noninjected ($\mathcal{G}_{\mathrm{Null}} \setminus \mathcal{G}_{\mathrm{spike}}$, $n = 5$) *Taco1, Fer1l5, Lst1, Tmem132e, Tada2b* | $n = 10$. Noninjected ($\mathcal{G}_{\mathrm{Null}} \setminus \mathcal{G}_{\mathrm{spike}}$, $n = 5$) *SKI, CORO1C, GDI1, SH3PXD2A, PPP3CB* |
| $\mathcal{G}_{\mathrm{spike}}$ injection set (SaLFC effect) | $n = 5$ *Mrc1, Atad3aos, Zbtb4, Tmem191c, Gm44710* | $n = 5$ *PRNP, PAM, PARVA, CD99, ZNF638* |

**Supplementary Table S4 | Gene sets for the platform-transfer cohorts.** Nested gene sets $\mathcal{G}_{\mathrm{spike}} \subset \mathcal{G}_{\mathrm{Null}} \subset \mathcal{G}_{\mathrm{SVG}}$ are shown for the Stereo-seq mouse embryo (lung versus neural tissue) and Visium HD colorectal cancer (normal epithelium versus stroma). $\mathcal{G}_{\mathrm{SVG}}$ is used for domain recovery, $\mathcal{G}_{\mathrm{Null}}$ for SaLFC testing, and $\mathcal{G}_{\mathrm{spike}}$ receives the imposed expression effect and contains the null genes with the largest baseline $|\Delta_g^{TR}|$. All sets were constructed using the same procedure as in DLPFC (Supplementary Note S1).
